# Cyclic tetra-adenylate triggers Cap1 membrane pore opening and depolarization during the type III CRISPR immune response

**DOI:** 10.1101/2025.11.13.688252

**Authors:** Puja Majumder, Clare W. Cahir, Christian F. Baca, Héloïse Carion, Cameron G. Roberts, Dinshaw J. Patel, Luciano A. Marraffini

**Author notes:** Present address: Bioscience and Biotechnology Department, Indian Institute of Technology, Kharagpur, WB 721302, India. Present address: Vertex Pharmaceuticals, 3215 Merryfield Row, San Diego, CA 92121, USA. These authors contributed equally (Puja Majumder, Clare W. Cahir).

## Abstract

Type III CRISPR-Cas systems synthesize cyclic oligoadenylates (cOAs) signaling molecules upon RNA-guided recognition of viral (phage) transcripts. cOAs bind to and activate CARF effectors, many of which contain transmembrane domains that cause membrane depolarization to prevent phage propagation. How ligand binding induces the activity of these effectors is poorly understood at the molecular level. Here we report the structure and function of the Cap1 effector, composed of a pair of transmembrane helices (TM1/2), a CARF-like (CARFL) domain and a domain of unknown function (DUF4579). *In vivo*, Cap1 activation results in membrane depolarization, a growth arrest of the bacterial host and the abrogation of the viral lytic cycle. Cryo-EM studies on apo– and cOA-bound states of Cap1 in glyco-diosgenin detergent revealed the formation of tetrameric complexes in both states, with one cOA molecule bound in a pocket composed of four CARFL domains. Ligand binding triggers conformational changes that widen an otherwise narrow transmembrane pore formed by the four TM1/2 domains. Our findings reveal at the atomic level the mechanism of pore opening and membrane depolarization mediated by cyclic nucleotide signaling during the type III CRISPR-Cas response.

## INTRODUCTION

Prokaryotic clustered regularly interspaced short palindromic repeat (CRISPR) loci and their associated genes (*cas*) provide RNA-guided adaptive immunity against invading genetic elements such as bacteriophages^1^ and plasmids^2^. A unique feature of these loci is their ability to acquire short sequences from the invading genomes that are stored as “spacers” in between repeats of the CRISPR locus^1^. The CRISPR array of repeats and spacers is transcribed and processed into short CRISPR RNAs (crRNAs) that are loaded into Cas ribonucleoprotein complexes and used as guides for the recognition of complementary nucleic acids^3,4^. Depending on the *cas* gene content, CRISPR-Cas systems can be classified into six types^5^. Type III systems encode the crRNA-guided Cas10 complex^3,6^, which is activated upon recognition of a complementary RNA molecule produced by the invader. Target finding triggers two enzymatic activities in the Cas10 subunit of the complex: ssDNA degradation by the HD domain^7,8^ and cyclic oligoadenylate (cOA) synthesis using ATP molecules as substrate by the Palm domain^9,10^. The ssDNase activity is presumed to directly attack the phage’s genome and is sufficient to provide immunity when the target transcript is expressed early in the viral lytic cycle^11^. In contrast, when phage RNA is recognized late during infection, type III CRISPR-Cas immunity requires the activation of diverse accessory effector proteins containing CARF (CRISPR-associated Rossman fold) domains^12–14^. CARF domains form a pocket that accommodates cOA and are linked to an effector domain that is activated upon ligand binding. CARF effectors orchestrate a broader antiviral response by triggering diverse activities that are toxic both for host growth and viral replication, such as DNA and RNA degradation^15–19^, toxic nucleotide accumulation^20,21^, NAD^+^ depletion^22^ and membrane disruption or depolarization^23,24^. In this way, CARF effectors turn type III CRISPR-Cas immunity into an abortive infection mechanism of defense that results in the sacrifice of the infected host to prevent the spread of the phage into the rest of the bacterial population^11^.

Of particular interest are CARF effectors harboring transmembrane domains that cause membrane depolarization, an activity that is difficult to study both *in vivo* owing to the membrane localization of the proteins, as well as *in vitro* due to the lack of biochemical assays. Previous studies of two of these effectors, Cam1^23^ and Csx23^24^, revealed that these effectors assemble into tetramers, bind cA_4_, and display potent anti-phage activity (through membrane depolarization in the case of Cam1). However, structural insight was limited to the isolated CARF domains resolved by x-ray crystallography, whereas the architecture of the transmembrane (TM) region was not experimentally determined and instead modeled using AlphaFold. The absence of a solved full-length structure precluded a mechanistic understanding of whether and how cA_4_ binding is coupled to a conformational change in the transmembrane domain, leaving the molecular details of the transition of these effectors from the resting to the active state uncertain.

Here we characterized Cap1 (CRISPR-associated pore 1), a CARF effector from the *Parafilimonas terrae* type III-A CRISPR-*cas* locus composed of two transmembrane helices, a CARF-like domain and a domain of unknown function (DUF4579). We demonstrate that Cap1 is a cyclic tetra-adenylate (cA₄)-activated effector that provides antiphage defense during type III CRISPR immunity via membrane depolarization. We solved cryo-EM structures of apo– and cA_4_-bound Cap1 in glyco-diosgenin (GDN) detergent and monitored pore formation within the membrane spanning segments of tetrameric scaffolds. We found that upon cA₄ binding to the CARF-like domain, the tetramer undergoes conformational changes that leads to pore opening. Our findings uncover structural details that establish a direct molecular link between cyclic nucleotide signaling and anti-phage defense through membrane depolarization by a CRISPR-associated membrane pore forming effector.

## RESULTS

### Cap1 induces membrane depolarization during the type III-A CRISPR-Cas response upon binding of cA_4_

*P. terrae* harbors a type III-A CRISPR-*cas* locus with a 26-spacer CRISPR array encoding for the Cas1/2 integrase responsible for the insertion of new spacer sequences^25–27^, the Cas10-Csm crRNA-guided complex^3,7^, the Cas6 endoribonuclease that processes the CRISPR array transcript into crRNAs^4,28^, a CARF effector and two ring nucleases that share structural homology to characterized ring nucleases Csx3^29^ (PDB 6VJG) and DUF1874 (PDB 6SCF) (Fig. S1A). Close inspection of the CARF effector protein sequence indicated that it is composed of a short N-terminal domain (N) followed by two transmembrane segments (TM1 and TM2) and a CARF-like domain (CARFL, see below) connected to DUF4579 through a linker (L) segment (Fig. 1A). Due to its function (see below), we renamed this protein “CRISPR-associated pore 1”, Cap1. In the past, we studied the function of CARF effectors *in vivo* in staphylococci, in the context of the *Staphylococcus epidermidis* RP62 type III-A CRISPR-*cas* locus. This locus is highly similar to the native CRISPR system that encodes Cap1 in *P. terrae* (Fig. S1A). We replaced the endogenous CARF effector *csm6*^30^ of the staphylococcal locus with *cap1*, and cloned the resulting locus into the staphylococcal vector pC194^31^, generating pCRISPR. We also introduced truncated versions of *cap1* with deletions of TM1 and TM2 region (*cap1-ΔTM*) or of the DUF4579 domain (*cap1-ΔDUF*) (Fig. S1B). As controls, we generated versions of pCRISPR with mutations that inactivate both activities of Cas10: D586A-D587A in the Palm domain (*cas10^Palm^*) to prevent the synthesis of cOA^9,10^ and H14A-D15A in the HD domain (*cas10*^HD^) to abrogate ssDNA degradation^8^ (Fig. S1B). In addition, a CRISPR locus without a targeting spacer (Δ*spc*) was used as a non-targeting control (Fig. S1B). pCRISPR plasmids were transformed into *Staphylococcus aureus* RN4220 cells^32^ containing pTarget, a second plasmid producing an anhydrotetracycline (aTc)-inducible transcript complementary to the crRNA carried by the Cas10-Csm complex encoded by pCRISPR^30^. In this two-plasmid system, addition of the inducer triggers type III-A CRISPR-Cas immunity and the synthesis of cOAs in the absence of phage infection (degradation of pTarget by the ssDNase activity of Cas10 is avoided using the *cas10*^HD^ allele in pCRISPR^30^). To determine if Cap1 is toxic during the type III-A CRISPR response, as is the case for most other CARF effectors^15,20,22,23,30^, we followed the growth of cultures carrying both plasmids through measurement of optical density at 600 nm (OD_600_), 360 minutes after addition of aTc. Cap1 and Cap1-ΔDUF, but not Cap1-ΔTM, prevented culture growth (Figs. 1B and S1C), in a manner that depended on the presence of a crRNA guide to recognize the aTc-inducible target transcript and on the synthesis of cOA by the Palm domain of Cas10 (Figs. 1B and S1C). To determine whether the failure of the cultures to grow after PtCap1 activation is caused by the death or lack of division of the affected cells, we measured the viability of the cells at different times after activation. We added aTc to staphylococcal liquid cultures and took aliquots over time, which we plated on solid media lacking the inducer to enumerate colony forming units (CFU) derived from cells that survive Cap1 activation. We found that the CFU count remained stable throughout the three hours after Cap1 activation (Fig. S1D), a result that demonstrates that cells in the culture are not dividing nor dying (the CFU did not increase nor decrease with time, respectively). Therefore, we conclude that Cap1 mediates a growth arrest of the bacterial population during the type III-A CRISPR-Cas response.

**Figure 1.**
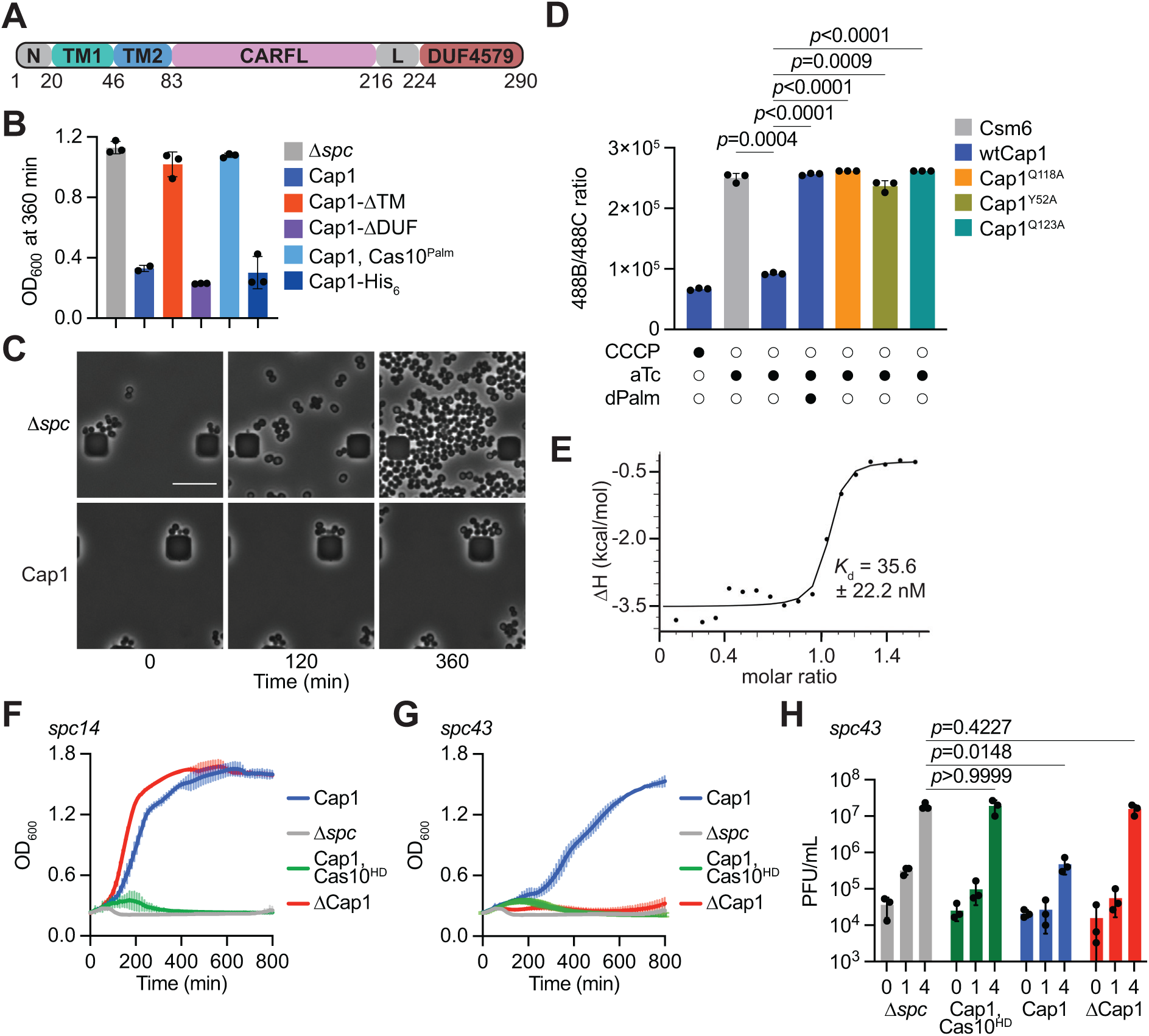
Cap1 mediates growth arrest upon activation of the type III CRISPR-Cas response. **(A)** The domain organization of Cap1 protein displaying the residue numbers for transmembrane segments (TM1, TM2), CARF-like domain (CARFL) and a domain of unknown function (DUF4579). **(B)** Growth of staphylococci carrying pTarget and different pCRISPR variants, measured as OD_600_ value at 360 minutes after the addition of aTc. Data show the mean of three biological replicates +/− s.e.m. **(C)** Images from live microscopy of staphylococci carrying pTarget and different pCRISPR variants, taken at different times after the addition of aTc. **(D)** Quantification of green/red fluorescence ratio (488B/488C channels) obtained using flow cytometry of staphylococci carrying pTarget and different pCRISPR variants stained with DiOC_2_(3) after induction of cA_4_ synthesis via aTc treatment of the cultures. The mean of three biological replicates +/− s.e.m are shown. *p* values were obtained with two-sided t-tests with Welch’s correction. **(E)** Binding of cA_4_ to the wild-type CARFL domain, measured by ITC; a *K*_d_ value of 35.6 +/− 22.2 nM was estimated from the curve. **(F)** Growth of staphylococci carrying different pCRISPR constructs targeting the *gp14* transcript produced by the ϕNM1γ6 phage, measured as OD_600_ after infection at MOI ∼5. The mean of three biological replicates +/− s.e.m are shown. **(G)** Same as in (F), but testing the immunity of pCRISPR constructs programmed to target the *gp43* transcript. **(H)** Number of plaque-forming units (PFU) in staphylococcal cultures harboring different pCRISPR constructs targeting the *gp43* transcript, at the indicated times after infection with ϕNM1γ6 at MOI ∼1. The mean of three biological replicates +/− s.e.m are shown. *p* values were obtained with two-sided t-tests with Welch’s correction.

To investigate the consequences of Cap1 activation at the cellular level, we followed the induced cultures using live microscopy. While the non-targeting control cells grew undisturbed over time, cells in which type III-A CRISPR immunity was triggered proliferated at a very low rate (Fig. 1C), a result consistent with Cap1 activation causing the arrest, but not the death, of staphylococci. Given that Cap1 is related to Cam1, a membrane-bound CARF effector that causes membrane depolarization^23^, we tested whether the growth arrest described above is caused by the same mechanism. We induced synthesis of the cA_4_ ligand using aTc, as in the toxicity assays experiments, and stained staphylococci with the membrane potential indicator dye 3,3’-dietyloxacarbocyanine iodide [DiOC_2_(3)]. The DiOC_2_(3) dye emits green and red fluorescence, with a shift towards red emission in the presence of higher membrane potentials^33^. Therefore, membrane potential can be measured independent of cell size by calculating the red/green fluorescence intensity ratios^33^. We used carbonyl cyanide *m*-chlorophenylhydrazone (CCCP), a common membrane potential disruptor, as a positive control^33^. As a negative control we used cells expressing Csm6, a CARF effector that does not affect the host membrane^30^, instead of Cap1. Flow cytometry analysis of the cells treated with DiOC_2_(3) showed that Cap1, but not Csm6, activation resulted in a low red/green fluorescence ratio only in the presence of a functional Palm domain capable of synthesizing cA_4_ (Figs. 1D and S1E).

Finally, since Cas10 can synthesize different cOA, cyclic tetra-(cA_4_), hexa-(cA_6_), and tri-adenylates (cA_3_)^9,10^, we decided to determine which one of these second messengers activates Cap1. We expressed a His-tagged version of the CARFL domain (residues 85-220; His_6_-Cap1-CARFL) in *Escherichia coli* cells and purified it using Ni^+2^ affinity chromatography followed by size exclusion chromatography (Figs. S1F-G). We used isothermal titration calorimetry (ITC) to determine the binding affinity of the purified protein for different cOA. We found that Cap1-CARFL-His_6_ binds cA_4_ with high affinity, with an estimated *K*_d_ value of 35.6 ± 22.2 nM (Fig. 1E). No interaction with cA_6_ nor cA_3_ was detected (Fig. S1H-I). In addition, upon addition of cA_4_, size-exclusion chromatography coupled with multi-angle static light scattering (SEC-MALS) analysis revealed a substantial shift towards a tetrameric molecular weight species (data not shown). Altogether, these results indicate that Cap1 is activated by the production of cA_4_ during the type III-A CRISPR-Cas response and causes membrane depolarization that leads to a growth arrest, but not death, of the host cell. Cap1 toxicity requires the transmembrane, but not the DUF4579, domain.

### Cap1 mediates immunity against phage infection

Next, we tested the role of Cap1 during the type III-A CRISPR-Cas response against phage infection. As a consequence of RNA targeting by the Cas10 complex^3,8^, this response depends on when the target transcript is produced during the viral lytic cycle^11,16,34^. CARF effectors are typically dispensable for defense when the target is recognized early after infection but absolutely essential when the phage transcript is detected by the Cas10 complex in late stages of infection^11,15,20,22,23^. We therefore programmed the CRISPR array of the pCRISPR plasmid with two spacers (*spc14* and *spc43*) producing crRNAs complementary to early– and late-expressed transcripts (*gp14* and *gp43*, respectively) from staphylococcal phage ϕNM1γ6. Each spacer was introduced into two different genetic backgrounds for the *cas10* gene: wild-type and cas10^HD^, carrying mutations in the nuclease active site (H14A, D15A) that prevent ssDNA cleavage^8^ (Fig. S1B). This mutation forces the type III-A immune response to rely exclusively on the CARF effector^11,30^, and therefore allows us to measure the ability of Cap1 to defend the host from infection. As a negative control we generated pCRISPR plasmids harboring *spc14* or *spc43* but without encoding a CARF effector (*Δcap1*, Fig. S1B). We infected staphylococci harboring the different pCRISPR plasmids with ϕNM1γ6 at a multiplicity of infection (MOI) of ∼5 and determined culture survival by measuring OD_600_. When programmed to target the early-expressed transcript (*gp14*), the type III-A CRISPR-Cas response required ssDNA degradation by the Cas10 complex, but not Cap1 (Fig. 1F). In contrast, when the target RNA is recognized late in the viral lytic cycle (*gp43*), Cap1 was necessary to support immunity and the nuclease activity of Cas10 was also required. Neither of these activities alone was sufficient to enable culture growth after infection (Fig. 1G). We corroborated these results by evaluating the effect of type III CRISPR-Cas immunity on phage propagation, enumerating plaque-forming units (PFU) on lawns of staphylococci carrying different pCRISPR(*spc43*) plasmids. We found a significant reduction in PFU only when hosts carried active Cap1 and Cas10, but not with either of these immune effectors alone (Fig. 1H). These results demonstrate that, as previously found for other CARF effectors^20,22,23^, the growth arrest mediated by Cap1 is required for a successful type III-A immune response that is activated late in the viral lytic cycle.

Since staphylococci are Gram-positive organisms, we decided to test its immune properies in a Gram-negative bacterium that would share a similar envelope composition as *P. terrae*, where Cap1 is encoded. This may be particularly important for Cap1 given its membrane localization, close to the surface of the host bacterium. We chose to test Cap1-mediated immunity in *E. coli*, against phage lambda infection. We were not able to clone the type III-A CRISPR-*cas* locus present in *P. terrae*, which harbors *cap1*, and therefore we used the previously characterized and highly similar *Serratia* sp. ATCC39006 type III-A CRISPR-Cas system^35,36^ (Fig. S1J). This locus harbors nine spacers and the cA_3_-activated SAVED effector NucC^36^ and was cloned into the low-copy-number plasmid pACYC184 (PMID 149110) (Fig. S1K). We then replaced the spacers by a single spacer that would target the lambda gene *O* transcript, involved in replication, to generate plasmid pCRISPR-Ss(NucC). We replaced *nucC* for *cap1* to construct pCRISPR-Ss(Cap1), and also removed this gene to generate a version of this plasmid that does not provide immunity as a negative control pCRISPR-Ss(*Δeff*). Cultures harboring these plasmids were infected with lambda phage at an MOI of ∼0.5 and their growth was followed by measuring OD_600_ (Fig. S1L). We found that, while cells without a cOA-activated effector collapsed upon infection, Cap1 expression was sufficient for cell survival, although it mediated a weaker growth than NucC, the native effector of this type III-A CRISPR-Cas system. We conclude that Cap1 can provide immunity to bacteria that possess both a Gram-positive and a Gram-negative envelope.

### Apo-Cap1 adopts a tetrameric architecture

To investigate the molecular mechanism behind Cap1’s toxicity, we expressed in *E. coli* a full-length version of Cap1 that harbors a C-terminal hexahistidyl tag (Cap1-His_6_). We confirmed that the addition of the tag did not alter the toxic properties of Cap1 (Figs. 1B and S1C). We then purified Cap1-His_6_ via Ni^+2^ affinity chromatography from the membrane fraction, in the presence of GDN (glycodiosgenin) detergent (Fig. S2A). First, we analyzed the oligomeric state of purified apo-Cap1 (in the absence of cOA ligands) using conjugate SEC-MALS, which rendered a somewhat broad peak indicative of tetramer formation (Fig. 2A, green arrow), along with the presence of some higher order species, mostly due to the mild aggregation propensity of the protein (Fig. 2A, black arrow). The peak fractions were collected to determine the atomic resolution structure of apo-Cap1 in GDN using cryo-EM. The collected images were separated in different representative 2D class averages (Fig. S2B). One class displayed a view from the cytosolic side that corroborated the tetrameric association suggested by SEC-MALS (Fig. S2B, image “*a*”). Another 2D class presented a side view of apo-Cap1, showing both the DUF and CARFL domains, as well as some of the TM helices surrounded by the detergent micelle (Fig. S2B, image “*b*”; Fig. S2C). The high resolution cryo-EM map of apo-Cap1 established a tetrameric arrangement composed of cytosolic DUF and CARFL domains connected to the TM1/2 membrane-spanning domain (Figs. 2B, S2C-D). In this structure, TM1 crosses the bacterial membrane starting with its N-terminal end in the cytosolic side and then takes a sharp turn at the extracellular side to allow the re-entry of TM2 into the membrane (Fig. 2B). Notably, in the tetrameric arrangement of apo-Cap1, this sharp turn leads to the formation an inner ring of four TM2 helices surrounded by an outer ring of four TM1 helices (Figs. 2B-C). Interestingly, we observed extra density protruding out from one of the TM2 helices inside the inner ring (Fig. S2E, black box). Upon building the protein model we found that this extra density most likely corresponds to an alternate conformation of the Y75 side chain of one of the four TM2 helices (Figs. S2F-G). We then evaluated the electrostatic potential of the inner ring using PyMOL and considered both conformations of the Y75 side chain. We found that in both cases the inner ring forms a closed pore with a positively charged cytosolic side that is surrounded by four negatively charged patches composed of residues T81, E82 and S83 in the TM2 segments (Figs. 2D, S2H). These residues are part of a short cytosolic extension of TM2 that separates the membrane from the cytosolic CARFL domains (Fig. 2B, red arrow).

**Figure 2.**
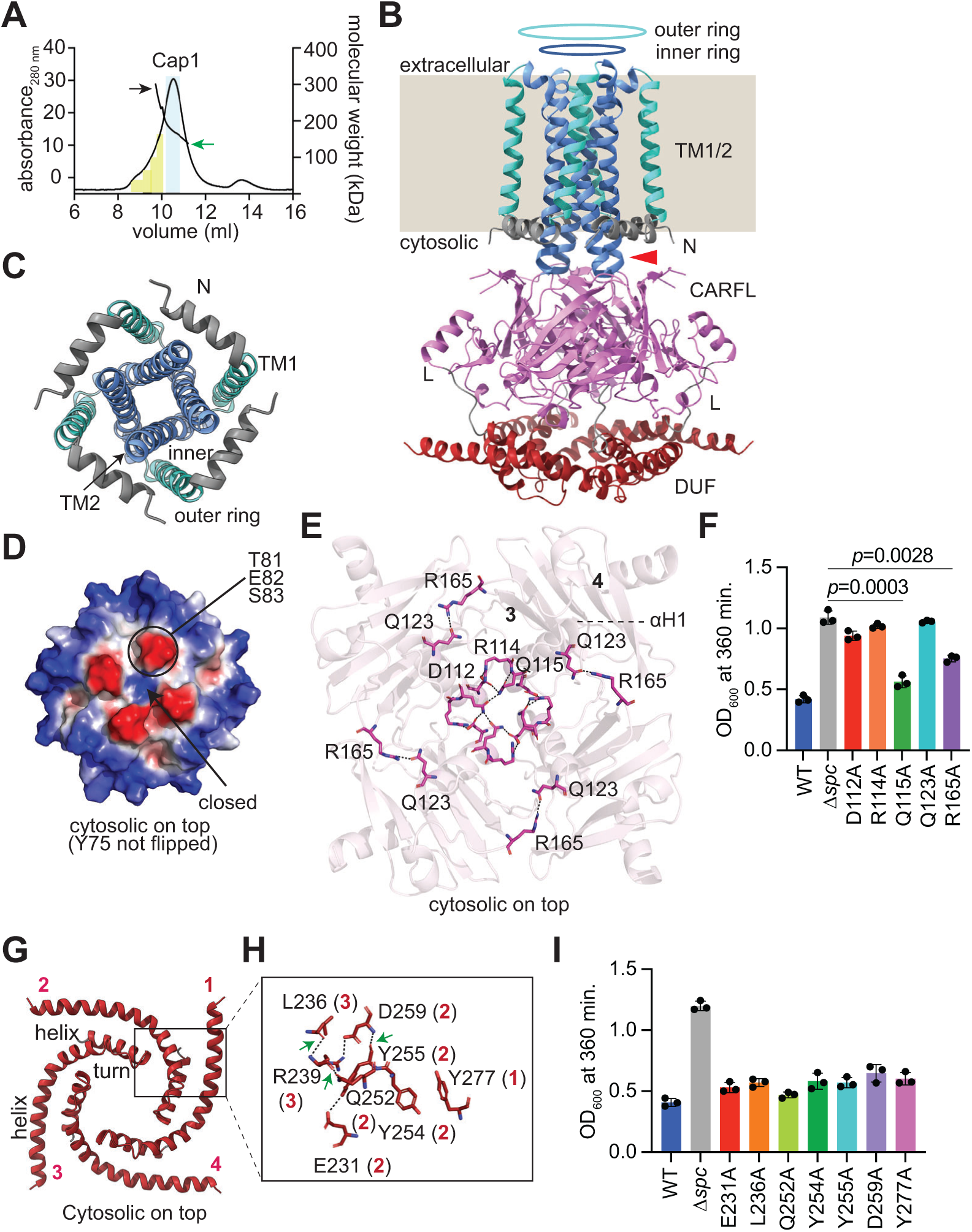
Apo-Cap1 tetramer structure. **(A)** SEC-MALS of apo-Cap1 protein. The major fraction elutes as a tetrameric form (green arrow, protein peak fraction is shown in light blue); other fractions contain species with higher oligomeric states (black arrow) that are generated mostly due to mild aggregation (marked in yellow) of the protein. **(B)** Ribbon diagram of the apo-Cap1 tetramer. TM1 and TM2 are displayed in green and blue and form outer and inner rings, respectively, that insert into the bacterial membrane (beige background, which replaces the density observed for the GDN detergent micelle). The cytosolic extension of TM2 is marked by a red arrow. The N-terminal segment (N, grey), CARFL domain (pink), the interconnecting linker (L, grey) and the DUF4579 domain (red) locate on the cytosolic side. **(C)** Cytosolic side view of the TM1 and TM2 domains of apo-Cap1 in green and blue, showing the formation of the outer and inner membrane rings, respectively. The N-terminal segment (N) is shown in grey. **(D)** Electrostatic surface representation of the cytosolic side view of the TM1 and TM2 domains, showing a closed pore. **(E)** Residues involved in CARFL tetramerization. D112, R114 and Q115 from αH1 of each monomer interact with each other at the center of the tetramer. R165 from the loop joining β strands 3 and 4 interacts with Q123 present in αH1. **(F)** Growth of staphylococci carrying pTarget and different pCRISPR variants harboring alanine substitutions of Cap1 residues shown in (E), measured as OD_600_ value at 360 minutes after the addition of aTc. Data show the mean of three biological replicates +/− s.e.m. *p* values were obtained with two-sided t-tests with Welch’s correction. **(G)** Four-fold arrangement of DUF4579 helix-turn-helix in the apo-Cap1 tetramer (each monomeric helix-turn-helix is numbered 1 to 4). The black box indicates the interaction site of adjacent helix-turn-helices. **(H)** Residues involved in helix-turn-helix interactions with the region marked in (G) with a black box. **(I)** Growth of staphylococci carrying pTarget and different pCRISPR variants harboring alanine substitutions of Cap1 residues shown in (H), measured as OD_600_ value at 360 minutes after the addition of aTc. The mean of three biological replicates +/− s.e.m are shown. *p* values were obtained with two-sided t-tests with Welch’s correction.

The CARF domain of apo-Cap1 (Cap1-CARF) assembles in a tetrameric conformation, with four αH1 helices at the center of the tetramer (Figs. 2E, S3A) that form a positively charged pocket with 4-fold symmetry (Fig. S3B). In most CARF effectors the ligand-binding pocket is formed by a dimer of CARF domains and accommodates the cOA second messenger^12–14^. In contrast, our cryo-EM data suggests that this effector captures its ligand using a tetrameric binding pocket, a highly unconventional arrangement. A more detailed structural comparison of Cap1-CARF with the canonical CARF domain of Cad1, a previously studied CARF effector (PDB 9C77)^20^, further differentiated both domains. 2D topology diagrams generated with Pro-origami^37^ showed that whereas the Cad1 CARF domain follows a classical Rossmann-fold (Fig. S3C), Cap1 CARFL adopts a more streamlined core architecture (Fig. S3D). Specifically, the central core of Cad1 CARF is formed by a 6-stranded β-sheet (β5, β3, β1, β7, β9 and β11) flanked by an elaborate array of six peripheral α-helices (α2, α4, α6, α8, α10 and α12). In contrast, Cap1 CARFL features a more compact, 6-stranded central β-sheet core (β7, β6, β2, β1, β5 and β4) supported by minimal helical decorations (only α3 and α8). These differences result in divergent spatial and electrostatic landscapes of the primary ligand-binding cleft of these domains. Due to these substantial differences, we decided to name the Cap1-CARF domain “CARF-like” (CARFL). The CARFL tetramer is stabilized by residues D112, R114 and Q115 located in αH1 that interact with each other and with equivalent residues in the αH1 of an adjacent monomer, as well as by the interaction between Q123 in αH1 and R165, situated in a loop joining β-strand 3 and 4 from a contiguous CARFL region (Fig. 2E). To assess the importance of tetramer formation and/or stability for Cap1 function, we generated single alanine substitutions of these residues and tested for the ability of Cap1 to cause toxicity upon activation of the type III-A CRISPR response. We found that while Q115A and R165A mutations did not alter Cap1-mediated toxicity, substitution of D112, R114 and Q123 for alanine abrogated growth arrest (Figs. 2F, S3E). In addition, cells expressing the Q123A mutant Cap1 did not undergo membrane depolarization upon activation of the protein (Fig. 1D). Given that Q123 interacts not only with R165 but also with the backbone of F132 (Fig. S3F), it is possible that the Q123–F132 interaction that remains in the R165A mutant is sufficient to stabilize the CARFL tetramer. Likewise, it is conceivable that in the Q115A mutant, the interactions between D112 and R114 (Fig. S3F) are able to maintain the tetrameric structure.

The cryo-EM structure of the DUF4579 of the apo-Cap1 revealed that it adopts a helix-turn-helix motif arranged in a tetraskelion-like shape in the tetramer (Fig. 2G). This shape is stabilized by interactions both within and between adjacent helix-turn-helices: E231, Q252, Y254, Y255 and D259 from monomer 2 interact with each other, with Y277 from monomer 1 and with L236 and R239 from monomer 3 (Fig. 2H). In contrast, we did not detect contacts between DUF4579 and CARFL residues. We tested the importance of the DUF4579 residues presumably involved in the interactions described above through the introduction of alanine substitutions. Interestingly, none of the mutations limited Cap1’s ability to induce growth arrest during the type III-A CRISPR-Cas response (Figs. 2I, S3G). This is in line with our previous result that showed a similar lack of effect on Cap1’s function for the deletion of the entire domain (Cap1-ΔDUF, Fig. 1B).

### The cA_4_-Cap1 tetramer binds a single cA_4_ molecule

To gain mechanistic insights into Cap1’s activation, we solved cryo-EM structures in the presence of cA_4_. The obtained images were separated into two distinct classes, with high and low resolution. The cryo-EM map of the first class at high (2.9 Å) resolution showed formation of a tetrameric complex, but without a visible DUF4579 (Figs. 3A, S4A). To ensure that this domain was not cleaved after cA_4_ addition, we compared the size exclusion chromatograms of apo– and cA_4_-bound Cap1. Since we obtained a very similar elution profile for both proteins (Fig. S4B), we concluded that the DUF4579 is still present after cA_4_ binding, but cannot be detected most likely due to high flexibility and/or structural heterogeneity.

**Figure 3.**
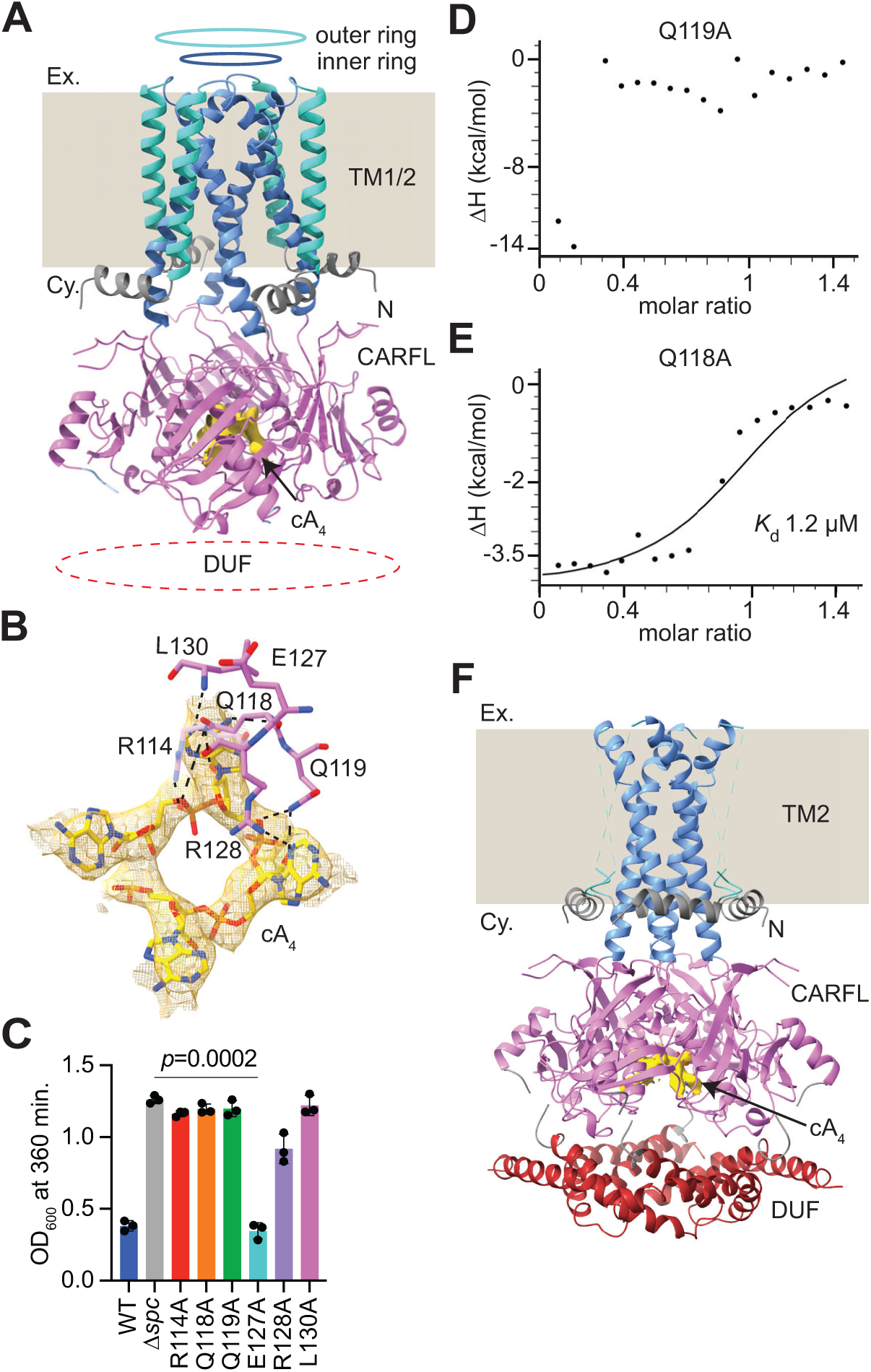
cA_4_-Cap1 structure. **(A)** Ribbon diagram of the cA_4_-Cap1 tetramer. TM1 and TM2 are displayed in green and blue and form outer and inner rings, respectively, that insert into the bacterial membrane (beige background, which replaces the density observed for the GDN detergent; “Ex.”, extracellular space, “Cy.” cytosol). The N-terminal segment (N, grey) and the CARFL domain (pink) locate on the cytosolic side. The density for the DUF4579 domain was not visible. **(B)** Residues of one CARFL monomer (pink sticks) that interact with cA_4_. The density for the cA_4_ molecule is shown in yellow mesh representation, at contour level ∼9 RMS from the C1 symmetry cA_4_-Cap1 map. **(C)** Growth of staphylococci carrying pTarget and different pCRISPR variants harboring alanine substitutions of Cap1 residues shown in (B), measured as OD_600_ value at 360 minutes after the addition of aTc. Data show the mean of three biological replicates +/− s.e.m. *p* values were obtained with two-sided t-tests with Welch’s correction. **(D)** Binding of cA_4_ to the CARFL^Q119A^ domain, measured by ITC. **(E)** Binding of cA_4_ to the CARFL^Q118A^ domain, measured by ITC; a *K*_d_ value of ∼1.2 µM was estimated from the curve. **(F)** Ribbon diagram of the cA_4_-Cap1 tetramer with a visible DUF4579 domain. TM2 helices are displayed in blue and form the inner ring that inserts into the bacterial membrane (beige background, which replaces the density observed for the GDN detergent; “Ex.”, extracellular space, “Cy.” cytosol). The N-terminal segment (N, grey) and the CARFL domain (pink) locate on the cytosolic side. The density for the outer ring of TM1 helices could not be modelled.

We observed one cA_4_ bound per tetramer, with the density corresponding to cA_4_ buried inside the CARFL tetrameric pocket, at the center of 4-fold symmetry, surrounded by αH1 helices (Figs. 3A, S4C-D). This is unusual for most cA_4_-CARF interactions, where cA_4_ binds to the pocket at the interface of a CARF dimer, being positioned on top of the two αH4 helices, as shown in the previously studied cA_4_-Cad1 structure (Fig. S4E)^20^. In addition, the adenine bases of the ligand are pointing upwards towards the cytosolic direction, rather than being positioned in the same plane as rest of the molecule (Figure S4F). We refined the cryo-EM map with C1 symmetry followed by C4 symmetry to improve the resolution. However, in both cases the CARFL domains were well resolved, and the map correspond to the cA_4_ molecule was similar (Figs. S4C-D); therefore, we used the C1 symmetry cA_4_-Cap1 map for further analysis of the ligand. In this structure, cA_4_ interacts with R114, Q118, Q119, E127, R128 and L130 residues (Fig. 3B). With the exception of E127A, alanine substitutions of these residues impaired the ability of Cap1 to mediate a growth arrest (Figs. 3C, S4G), with the mutation Q118A also impairing membrane depolarization (Fig. 1D). We also purified Cap1-CARFL-His_6_ domains carrying the Q118A and Q119A mutations (Figs. S1G, S4H-I) and performed ITC experiments to measure cA_4_ binding directly. We observed severe reductions in ligand binding, with the Q119A substitution completely abrogating cA_4_ interaction (Fig. 3D) and the Q118A mutation decreasing binding by more than two orders of magnitude (*K*_d_ ∼ 1.2 µM; Fig. 3E).

The cryo-EM map of the low resolution class (3.6 Å) showed the DUF4579, the CARFL and TM2 domains, however the TM1 helices were not resolved (Figs. 3F, S4J). The DUF4579 adopted a similar tetrameric conformation as in apo-Cap1 (Fig. S4K). In contrast, the cA_4_ binding site and the arrangement of cA_4_ in this structure were similar to those observed in the high resolution cA_4_-Cap1 complex, with comparable interacting residues (Fig. S4L). Although this mode of ligand binding (one cA_4_ bound per tetramer) was not observed in any of the previously solved structures of CARF/SAVED/Csx3 domains, a similar arrangement was observed for the isolated cytosolic domain of Csx23^24^, a cA_4_-activated type III CRISPR-Cas transmembrane effector (Figs. S4M-N). Altogether these results demonstrate that Cap1 binds cA_4_ molecules with an unconventional architecture in which four CARFL domains are required to accommodate a single ligand molecule.

### Phylogenetic analysis of PtCap1 homologs reveals a conserved cA_4_ binding mode by the CARFL domain

The unique structures we obtained of Cap1’s CARFL domain, with and without its ligand, prompted us to examine in more detail the phylogenetic distribution of this domain as well as its sequence and structural conservation across phyla. PSI-BLAST^38^ using PtCap1 sequence identified over 100 homologs (Supplementary File S1 and Fig. S5A), most of which encode both transmembrane helices and CARFL domains (96% and 99%, respectively), and 57% were located within type III CRISPR loci, although this is likely an underestimation given the presence of Cap1 homologs within incomplete genomes. Cap1 homologs were detected in at least 15 bacterial phyla, highlighting their broad distribution (Fig. S5A). Sequence alignment of the CARFL domains revealed a conserved motif, R(X_3_)QQ(X_10_)L, contained within the αH1 helix and capping loop characteristic of the PtCap1 structure (Figs. S3D and S5B). To perform structure-base analysis, we used FoldSeek^39^ to obtain 29 homologs of the monomeric structure of PtCap1, which we subjected to FoldMason^40^ analysis to generate a structural tree (Fig. 4A). Cap1 structures were found in at least eight bacterial phyla and were separated into two major clades: one consisting primarily of transmembrane helices fused to CARFL domains and another comprising longer proteins with additional domains. From these, C-terminal DUF4579 fusions were rare and largely confined to a small subclade. Three representatives of this clade were modelled using AlphaFold3^41^ (AF3), which predicted a tetrameric membrane-spanning pore architecture similar to the cryoEM structure of PtCap1 (Fig. 4B). Moreover, the αH1 helix and capping loop that contain the R(X_3_)QQ(X_10_)L cA_4_-binding motif showed good structural superposition, including the key conserved residues (Fig. 4C). We performed a similar analysis for Cad1^42^ homologs that contain canonical CARF domains (Fig. S3C). We used Foldseek to retrieve 15 homologs with highly similar structures, whose sequences were used to make a phylogenetic tree (Fig. S6A). The Cad1 homologs belonged to five different phyla and one representative of each was used to obtain an amino acid sequence alignment (Fig. S6B), which revealed conservation of residues S12, T35, K39, D67 and K104, i.e., different than the residues conserved in CARFL domains (Fig. S5B). A structural alignment of three selected homologs (Fig. S6C) showed that these residues are disseminated across multiple secondary elements, a result that contrasts with the compact and localized distribution of conserved residues in Cap1-CARFL. Altogether, these findings demonstrate that CARFL domains evolved a structural solution for cOA binding divergent from canonical CARF domains that is preserved across a wide variety of prokaryotes.

**Figure 4.**
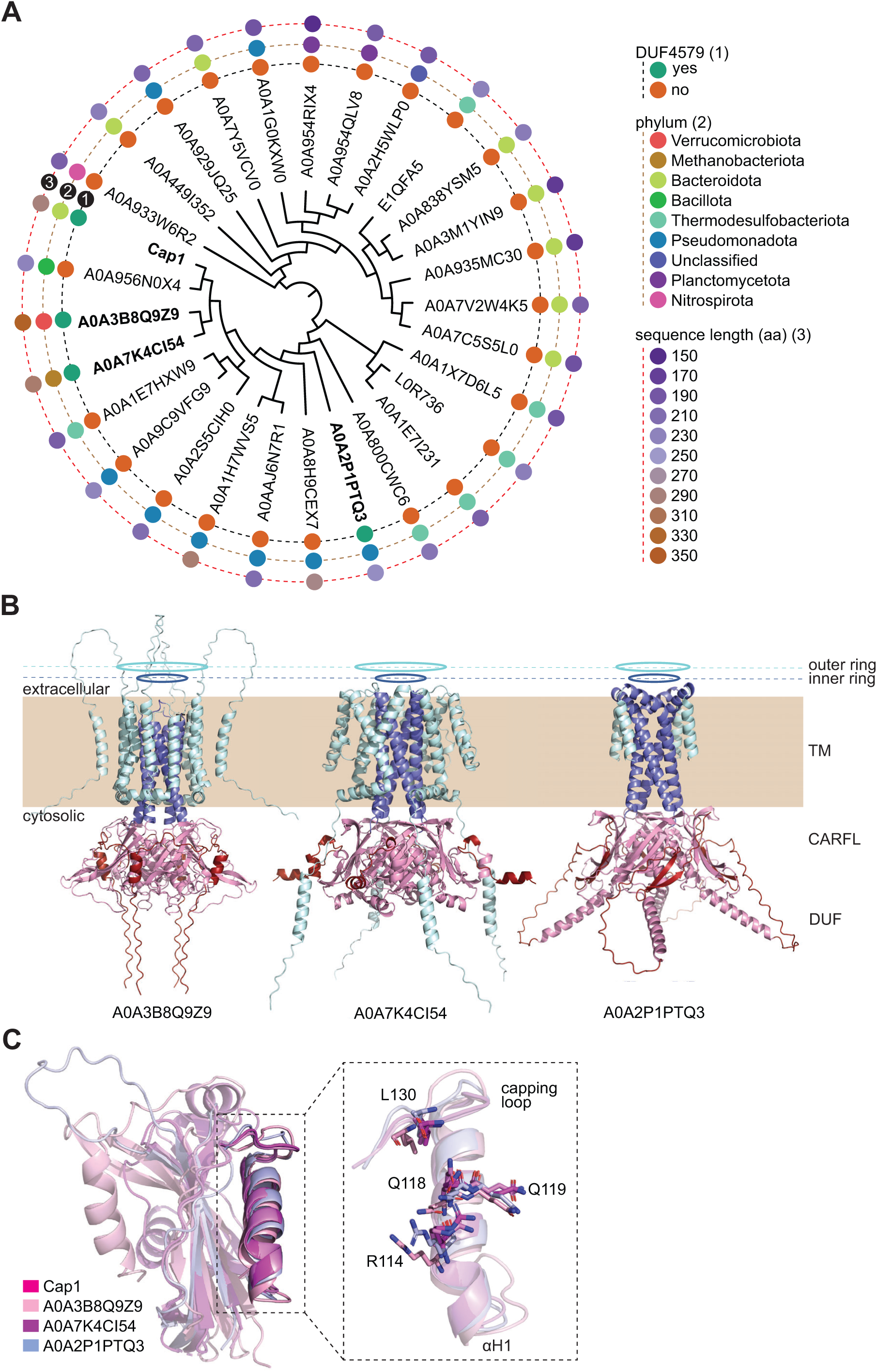
Phylogenetic analysis of Cap1 homologs. **(A)** Phylogenetic tree of PtCap1 monomer structural homologs. Structural homologs were found using FoldSeek (identified by their accession numbers); the top 29 hits were aligned with FoldMason to construct the tree. Three properties of each homolog are indicated: the presence of DUF4579 (1), the host phylum (2) and the sequence length (3). **(B)** AlphaFold3 tetrameric models of the PtCap1 structural homologs shown in bold in (A). **(C)** Structural alignment of AlphaFold models of monomeric CARFL domains shown in (B) and the cryo-EM structure of monomeric apo Cap1 CARFL, highlighting the αH1 helix and the capping loop. The inset shows a detailed view of the superposition of these structural elements, which contain the conserved cA_4_ binding motif residues R(X_3_)QQ(X_10_)L.

### cA_4_ binding opens Cap1’s transmembrane pore

Structural alignment of the apo– and cA_4_-bound Cap1 tetramers revealed ligand-induced conformational changes in the TM1/2 and CARFL domains (Fig. 5A), as well the loss of resolution of the DUF4579 domain. The most significant of these changes is a 6 Å outward shift of the inner TM2 helices (as viewed from the cytosolic side, Fig. 5B, blue arrows) that leads to the opening of the pore formed by the TM1/2 domains. We used Mole2.5 software^43^ to analyze the pore in the resting and cA_4_-activated (Fig. 5C) states. Upon activation, the cytosolic gate of the pore expands markedly, with the pore radius increasing from ∼3 Å in the resting state to ∼7–8 Å in the cA_4_-bound state (Fig. 5D). In the inactive state, the narrowest part of the pore is approximately at midlength, where residue T64 is located, with a radius of 1 Å (Fig. 5D). A similar radial constriction is observed in the apo-Cap1 structure with Y75 flipped-in (Fig. S7A), where the inward rotation of the Y75 side chain partially occludes the pore, shortening its length by approximately 1 Å relative to the inactive state structure where Y75 is flipped out (Figs. 5C and S7A).

**Figure 5.**
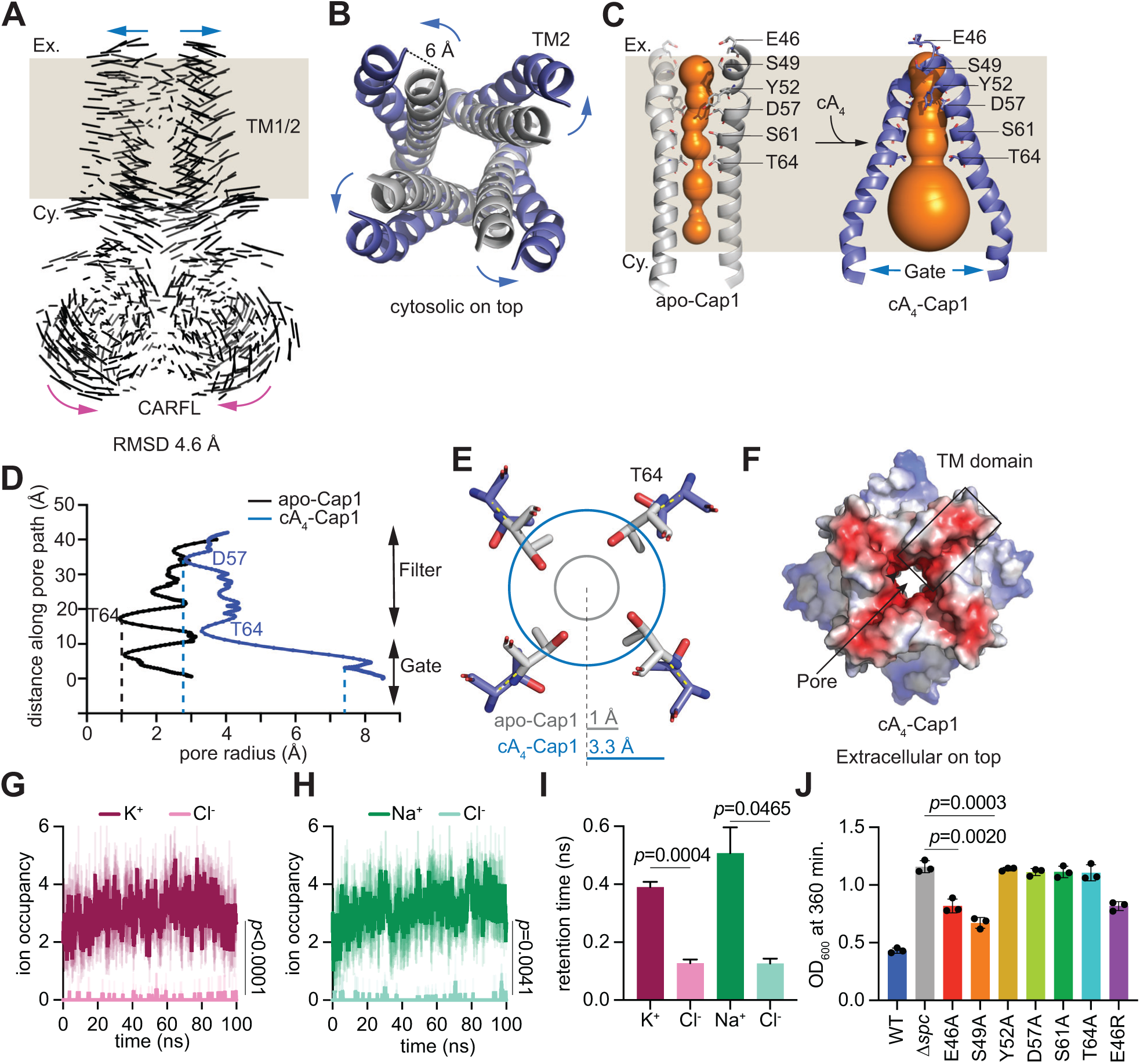
Pore opening upon cA_4_ binding causes membrane depolarization. **(A)** Superposition of apo– and cA_4-_-bound Cap1 (without visible DUF4579) structures shows that ligand binding promotes an outward movement of the TM1/2 helices (blue arrow) and an inward movement of the CARFL domain (pink arrow). The bacterial membrane is represented by the beige background; “Ex.”, extracellular space, “Cy.”, cytosol. The overall Cα RMSD between structures is 4.6 Å. **(B)** Cytosolic view of the outward movement (blue arrows) of TM1 inner ring helices. Helices in apo-Cap1 and cA_4_-Cap1 are shown in grey and blue, respectively. **(C)** Analysis of the pore formed by the TM1/2 domain using Mole2.5 software, represented in orange inside the TM2 helices. Apo-Cap1 structure shown has the Y75 side chain in a not flipped alignment. The bacterial membrane is represented by the beige background; “Ex.”, extracellular space, “Cy.”, cytosol. The narrowest part of the pore is formed almost at the middle of TM2 and is surrounded by four T64 residues. Upon cA_4_ binding, the pore opens towards the cytosolic side (in the structure without visible DUF4579). **(D)** Pore radius (calculated with Mole2.5 software) across the pore length for apo– and cA_4_-bound forms of Cap1, showing the position of key pore-lining residues. The filter (extracellular side) and gate (cytoplasmid side) regions of the pore are shown. **(E)** Cross-sectional view of the T64 side chains within the pore in the apo– and cA_4_-bound forms of Cap1 in grey and blue sticks, respectively. The closed (grey) and opened (blue) pore circumferences and radii at the cross-section, calculated with Mole2.5 software, are indicated. **(F)** Electrostatic surface representation of the extracellular side view of the TM1/2 domain, showing an opened pore with an electronegative lining. **(G)** The ion occupancy within the opened Cap1 pore in the presence of KCl, over 100 ns molecular dynamic simulations, for three independent replicates is displayed. **(H)** Same as **(G)** but simulating the presence of NaCl. *p* values were obtained using two-sided paired *t*-tests. (**I**) Mean ion residence (retention) times for K⁺, Na⁺, and Cl⁻ ions within the Cap1 pore, calculated from three independent molecular dynamics simulations in the presence of KCl and NaCl. Error bars represent mean ± s.e.m across replicates. *p* values were obtained with two-sided *t*-tests with Welch’s correction. **(J)** Growth of staphylococci carrying pTarget and different pCRISPR variants harboring alanine substitutions of Cap1 residues shown in (H) (and also E46R), measured as OD_600_ value at 360 minutes after the addition of aTc. The mean of three biological replicates +/− s.e.m are shown. *p* values were obtained with two-sided t-tests with Welch’s correction.

In the cA_4_-activated state the pore is narrower at the extracellular side and at the position of residue T64, where the radius is ∼3.3 Å, it opens towards the cytosolic region (Figs. 5C-E). Residues lining the narrow stretch are predominantly negatively charged or polar, resulting in an extracellular pore entrance that may interact with cations (Figs. 5C,F). The minimum radius of the pore is ∼2.8 Å at residue D57 (Fig. 5D), which is compatible with the passing of sodium and potassium ions. To determine if this is possible, we performed molecular dynamic simulations of cA_4_-bound Cap1 embedded in a lipid bilayer in the presence of KCl and NaCl salts and found that the cation, but not the anion, entered the pore (Fig. S7B). Furthermore, pore-lining residues E46, S49, Y52, D57, S61, and T64 were observed to be in close proximity with potassium and sodium ions within the pore (Fig. S7C). Quantitative analysis of ion trajectory in these simulations showed that K^+^ (Fig. 5G) and Na^+^ (Fig. 5H) have a significantly higher steady-state occupancy than Cl^-^ counterions, consistent with an electrostatically favorable selectivity filter formed by negatively charged or polar pore-lining residues that promotes cation coordination. Similarly, ion dwell time analysis showed prolonged retention time for K⁺ and Na⁺ than for Cl^-^ (Fig. 5I), a measurement that supports preferential cation occupancy within the Cap1 pore.

To verify the importance of pore lining residues we introduced alanine substitutions and found that Y52A, D57A, S61A and T64A completely impaired Cap1’s-mediated toxicity in staphylococci, with E46A and S49A displaying a less drastic effect (Figs. 5J, S7D). We also introduced a positive charge in position 46 (E46R mutation), however this substitution did not have a more profound effect on cell growth (Figs. 5J, S7E). In addition, the Y52A mutation abrogated membrane depolarization (Fig. 1D). Although there is the chance that these mutations (or any other mutation tested in this study) could affect the folding and/or membrane localization of Cap1 tetramer, the results emphasize the importance of the pore lining residues located in the TM2 domain for Cap1 activity and support a model that membrane depolarization occurs by the passage of cations through the pore.

Regarding conformational changes detected for the other domains of Cap1 upon ligand binding, we found that the αH1 helices of the CARFL domain showed an inward shift of 3.5 Å (Figs. 5A, S7F, magenta arrows). This is an opposite movement to that experienced by the TM1/2 domains and leads to the formation of a compact cA_4_ binding pocket. A major difference between apo Cap1 and the higher resolution structure of cA_4_– bound Cap1 structures was the absence of a defined DUF4579 in the latter. We used AF3 to corroborate this observation. The best four AF3 models of apo-Cap1 (interface predicted template modeling score, ipTM, 0.81; predicted template modeling score, pTM, 0.81) yielded similar structures to the one obtained by cryo-EM, with a good alignment of the four predicted DUF4579 structures (Fig. S7G). Superposition of the cryo-EM and top AF3 DUF4579 structures (red and silver, Fig. S7H) showed a good agreement between the two, with a Cα RMSD of 0.5 Å. In contrast, the best four AF3 models for the cA_4_-Cap1 (ipTM 0.71; and pTM 0.72) tetramer displayed a good alignment through the TM1/2 and CARFL domains; however, the DUF7549 remained completely miss-aligned and highly heterogeneous (Fig. S7I). Therefore, AF3 modeling supports our hypothesis based on cryo-EM results that cA_4_ binding generates a conformational change that leads to a disordered DUF4579.

Finally, superposition of the cA_4_-bound Cap1 structure with a visible DUF4579 to the apo form revealed minimal conformational changes in the CARFL and TM2 domains, with a Cα RMSD of 1.3 Å (Fig. S7J). This is a better structural alignment for these domains than that between both cA_4_-Cap1 structures (with and without visible DUF4579 domain), which has a larger Cα RMSD of 2.4 Å (Fig. S7K). This suggests that in the presence of the DUF4579 structure, the CARFL region bound to cA_4_ is not fully compacted and the TM2 domain retains a conformation more similar to that of the apo form (Fig. S7L), with a constricted pore (Fig. S7M). Based on these results, we speculate that the ordered DUF4579 functions as a structural gatekeeper, stabilizing the pore in a constricted configuration. We hypothesize thar full pore opening likely requires additional conformational transitions upon ligand binding of the DUF4579 assembly that remain unresolved, potentially due to high flexibility.

## DISCUSSION

In this study we elucidated the structure and mechanism of action of Cap1, a type III CARF effector. We show that Cap1 adopts a tetrameric architecture, with eight hydrophobic helices forming a pore that spans the bacterial membrane and four CARF– like (CARFL) domains that form a pocket for a single cA_4_ molecule. In the absence of ligand, the pore is closed but opens upon binding of the second messenger produced after the RNA-guided recognition of phage infection that triggers the type III-A CRISPR-Cas response. Cap1 activation leads to membrane depolarization, presumably due to the passage of ions across the open pore. The nature of the transported ions remains unknown, although the negative charges that line the pore, as well as our molecular simulations, suggest a model consistent with a preference for cation occupancy. Membrane depolarization is toxic to the host, which stops growing to prevent phage propagation.

Previous work covered three transmembrane effectors that participate in the CRISPR– Cas immune response. VanderWal *et al*. reported that Csx28, which is associated with type VI-B CRISPR-Cas systems^44^, displays an octameric configuration that forms a pore across the bacterial membrane. Csx28 is required for anti-phage defense, presumably through induction on membrane perturbations, but since type VI systems do not synthesize second messengers, it is not clear how it is activated and whether Csx28 undergoes conformational changes. Baca *et al*. described Cam1, a CARF effector linked to a short hydrophobic helix that localizes to the membrane^23^. Cam1 is part of a type III-A CRISPR-Cas system and is activated by cA_4_ upon crRNA-guided recognition of the infecting phage transcripts to cause membrane depolarization and prevent the viral lytic cycle. Although AF3 models predict the formation of a tetrameric pore across the membrane, the structure of full-length Cam1 in either apo– or cA_4_-bound states is not available. Finally, Grüschow *et al*. characterized Csx23, an effector associated with a type III-B CRISPR-Cas system with a unique cA_4_ binding domain fused to a short transmembrane domain whose activation is toxic to the host^24^. Using Pulse Dipolar Electron Paramagnetic Resonance Spectroscopy (PDS) to measure conformational changes, it was shown that cA_4_ binding to the purified full-length protein caused changes in the PDS signal that are consistent with structural heterogeneity in the transmembrane region. Outside of the CRISPR-Cas response, a group of effectors from cyclic oligonucleotide-based antiphage signaling systems (CBASS) contain transmembrane helices (Cap14, Cap15 and Cap16) that bind cyclic nucleotide second messengers to oligomerize and disrupt the membrane of infected cells^45^. While Cap1 is similar to these immune effectors in that they contain transmembrane domains and cause membrane disruptions, their characterization lacked structural data to precisely describe the nature of the changes triggered by ligand binding. In contrast to these previous studies, we demonstrate with atomic resolution how cA_4_ binding to the CARFL domain of Cap1 results in an outward movement of the transmembrane helices that opens a pore, causing membrane depolarization to compromise the infected host and thus prevent the propagation of the invading virus.

Cryo-EM analysis uncovered that Cap1’s CARFL domain diverges from conventional CARF domains both in protein fold and oligomeric assembly. While most of these structures typically form dimers with two-fold symmetry and bind one cA_4_ molecule at the dimer interface, CARFL assembles into a tetramer with four-fold symmetry and binds a single cA_4_ molecule at the central tetrameric interface – revealing a fundamentally different mode of ligand recognition. Notably, the central pore traverses not only the TM segments, but also the CARFL domains including the central hole of the bound cA_4_ second messenger. The previously characterized C-terminal cA_4_-binding domain (CTD) of Csx23^24^ and the CARFL domain of Cap1 adopt a similar tetrameric arrangement that binds a single cA_4_ molecule. Although both share a similar oligomeric arrangement, the Csx23 CTD comprises two antiparallel β-strands linked to a third by two α-helices, whereas the Cap1 CARFL domain forms a mixed β-sheet of two parallel and three antiparallel strands flanked by two α-helices, yielding a structurally distinct and bulkier domain architecture. The uncovering of these two structures that significantly diverge from canonical CARF domains suggests the possibility that many more yet uncharacterized cOA-binding domains exist.

Phylogenetic analyses revealed a broad distribution of Cap1 homologs containing CARFL domains across diverse bacterial hosts. This result underscores both the conservation of transmembrane depolarization as an antiviral strategy as well as the evolutionary flexibility of CARF domains to bind their ligands. It is interesting to speculate about the existence of additional CARF domains with unconventional architectures to accommodate other ligands, different than the reported cOAs and SAM– AMP molecules^46^. The evolution of non-canonical CARF domains most likely responds to the raise of phages that express type III CRISPR inhibitors to sequester the second messengers deployed during the immune response^47^. More broadly, alternative CARF folds may also have evolved binding flexibility to receive signaling molecules from other systems that utilize nucleotide signaling such as CBASS, Thoeris, and Pycsar^48,49^.

Cap1 also contains a domain of unknown function, DUF4579, that adopts a tetrameric fold, with each protomer forming a helix-turn-helix motif. This domain is connected through a linker to CARFL and caps it from the bottom. There are no detectable side chain interactions between CARFL and DUF4579 in the apo-Cap1 cryo-EM structure. Intriguingly, DUF4579 appears to be disordered in the cA_4_-bound state of Cap1, suggesting increased flexibility. Consistently, AF3 predictions show high structural heterogeneity for this domain when cA_4_ is included during modeling, reinforcing the idea that cA_4_ binding induces conformational variability or destabilization of this region.

Neither mutations that disrupt the tetrameric fold of this domain nor its complete deletion affected Cap1’s toxicity. These observations made us hypothesize that the DUF4579 may either be dispensable for Cap1’s function or may function as a structural gatekeeper, stabilizing the closed pore conformation of the apo state. The second scenario is supported by our finding of a minor cA_4_-bound cryo-EM class in which the DUF4579 remains ordered and the membrane pore retains a constricted architecture closely resembling the apo state. This configuration could represent an intermediate of the activation pathway, in which cA_4_ binding has occurred but neither conformational changes that increase in the flexibility or disorder in DUF4579 nor full pore opening has yet been achieved. *In vivo*, this domain is dispensable for Cap1 function, at least under our experimental conditions, but it is conceivable that it can play a specific role in its native host or when bacteria are growing in different media. Future studies hopefully will elucidate the importance of DUF4579 and other unknown domains linked to type III CRISPR-CARF effectors.

## METHODS

### Bacterial growth

*Staphylococcus aureus* strain RN4420^32^ was grown in brain heart infusion (BHI) medium at 37 °C, supplemented with chloramphenicol at 10 µg ml^-1^ for maintaining pCRISPR and erythromycin at 10 µg ml^-1^ for maintaining pTarget. 5 µM CaCl_2_ was supplemented in phage experiments unless indicated otherwise.

### Plasmid construction

The plasmids, oligonucleotides and cloning strategies for generating the plasmids used in this study are detailed in Supplementary S2. Cap1 coding sequence, NCBI GenBank reference sequence WP_090655733.1 from *Parafilimonas terrae* bacterium, was synthesized by Genewiz.

### PtCap1 sequence homology search

To determine how widespread Cap1 homologs are, we performed a PSI-BLAST search using PtCap1’s protein sequence. 134 homologs were detected and further analyzed. Homologs were then aligned using Clustal Omega server using default parameters^1^.

The resulting homolog alignment was used as an input to construct a phylogenetic tree with iqtree3^2^ with the –m TEST –bb 1000 flags. The resulting tree file was used as an input for iTOL^3^ for visualization. To determine and annotate whether homologs contain a CARFL domain, a hmm model was created for this domain using hmmbuild from HMMER v3.4^4^ with PtCap1 residues 83-216 as an input. The resulting hmm was searched across the homologs with hmmsearch using default parameters and an E– value cutoff of 1e-3. To predict and annotate whether homologs possess transmembrane helices, each corresponding protein sequence was scanned with deepTMHMM v1.0.57^5^. Finally, to determine whether host organisms for each homolog possess type III CRISPR systems, associated host nucleotide accessions were downloaded and analyzed with DefenseFinder v1.3.0^6^. If the resulting ‘gene’ file for each nucleotide fasta possessed a gene annotated as type III CRISPR-associated the homolog was annotated as positive for possessing a type III CRISPR system in the host genome.

### PtCap1 structural homology search

To further investigate the distribution and evolutionary relationship of Cap1 folds, a structural homology search using FoldMason^7^ from FoldSeek^8^ hits with the AlphaFold database^9^ model for PtCap1 (UniProt ID: A0A1I5TJE5). 29 FoldSeek structural homologs in the AFDB50 database with E-values less than 1e-3 were used to construct a structural alignment and tree using FoldMason with default parameters. The resulting tree was visualized with iTOL^3^. To annotate whether a C-terminal DUF was possessed by each structural homolog, each homolog was visually analyzed via an alignment to PtCap1’s monomeric AFDB model.

### Cad1 CARF and Cap1 CARFL 2D topology analysis

Two-dimensional topology diagrams of CARF domains from Cad1 (PDB ID: 9C77) and the CARFL domain of Cap1 cryo-EM structure were generated using Pro-origami^37^ to visualize secondary structure connectivity and folding architecture.

### Cad1 CARF structural homology search and dataset curation

The CARF domain of Cad1 (PDB ID: 9C77) was used as a query for structural similarity searches against the AFDB50 Database. Searches were performed using Foldseek^39^.

The top 15 hits were selected for downstream analysis (E value 2.5e-17 to 8.20e-10). To capture phylogenetic diversity, one representative sequence per bacterial phylum was randomly selected from the 15 structural homologs.

### Sequence conservation and motif mapping

Multiple sequence alignments were performed to map conservation patterns within the predicted cA₄/cA₆-binding pocket of CARF domain of Cad1. Conserved residues were identified and manually mapped onto structural models to assess their spatial distribution across homologs. Similar studies were performed with Cap1 CARFL homologs.

### Structural alignment of ligand-binding sites

Three representative Cad1 CARF homolog structures were selected and structurally aligned using Pymol software. The relative positions of conserved residues within the ligand-binding pocket were visualized to assess conservation of spatial organization across homologs. Similar studies were performed with Cap1 CARFL homologs.

### Growth curves

For *in vivo* Cap1 toxicity induction, biological replicates of RN4220 overnight cultures containing pTarget and pCRISPR are diluted 1:100, outgrown for about 75 min and normalized for optical density. Cells are then seeded in a 96-well plate. To induce targeting, 125 ng ml^-1^ of aTc is added to the appropriate wells. Absorbance at 600 nm is then measured every 10 min by a microplate reader (TECAN Infinite m200 PRO).

For *in vivo* antiphage immunity, cells containing various pCRISPRs were launched in triplicate overnight, diluted 1:100, outgrown for about 75 min and normalized for optical density. Cells were supplemented with 5 mM CaCl_2_ and seeded into a 96-well plate. ΦNM1γ6 was added at the specified MOIs, and optical density measurements were taken every 10 min.

For *in vivo* antiphage immunity in *E. coli*, cells containing various pCRISPR-Ss plasmids were launched in triplicate overnight, diluted 1:100, outgrown for about 90 min and normalized for optical density. Cells were supplemented with 1 mM MgCl_2_ and 1 mM CaCl_2_ and seeded into a 96-well plate. Lambda phage was added at the specificed MOIs, and optical density measurements were taken every 10 min.

### Cap1 viability assay

To measure the effect of Cap1 activity on *S. aureus* viability over time, colonies of *S. aureus* containing pTarget and the specified pCRISPR were launched in liquid culture overnight triplicate. The next day, cells were diluted 1:100 and grown out for about 1 hour and normalized for optical density. One aliquot was taken from each culture, and then aTc was added to induce CRISPR targeting and Cap1 activity. At each time point, cell aliquots were removed, centrifuged and resuspended in medium lacking aTc. All viable cells should grow on solid agar plates, but only targeting escapers (cells that recover owing to mutations in pTarget or pCRISPR) should form colony-forming units on plates with aTc.

### Quantification of phage plaques

To obtain plaque-forming unit (PFU) counts over time from cultures infected with phage, *S. aureus* cultures containing various pCRISPRs were launched overnight, diluted 1:100 and outgrown for about one hour. Cells in media supplemented with 5 mM CaCl_2_ were then infected with phage ΦNM1γ6 at an MOI of approximately 5, and an aliquot, filtered through a 0.45 µm membrane (Pall Acrodisc), was taken shortly after to obtain plaque-forming units at time 0. The cultures were then incubated further, with aliquots taken at one and four hours. For phage plaquing assays, indicated phage stocks were plaqued on lawns of *S. aureus* containing the indicated constructs, or cells lacking an introduced pCRISPR plasmid to measure PFU changes over time, with 10-fold serial dilutions for every spot in a lane.

### Flow cytometry

For our membrane depolarization studies, colonies of *S. aureus* containing pTarget and the specified pCRISPR were launched in liquid culture overnight. The next day, cells were diluted 1:100 and grown out for about 75 min and normalized to 10^7^ cells ml^−1^ in PBS. These cultures were then split into different subcultures and treated with either 125 ng ml^−1^ aTc or 1.7 μM CCCP (Thermo Fisher). These subcultures were incubated in shaking conditions at 37 °C for 30 min followed by addition of 15 μM DiOC_2_ (3) (Thermo Fisher) and incubation at room temperature for 5 min. Cells were then analyzed on a BD LSR-Fortessa (BD Biosciences) using FACSDiva Software version 8.0.1 with 100,000 post-gating events recorded for each sample. Red/green ratios were calculated using the Ratio Tab of the FACSDiva Software. Each cell’s red/green ratio was calculated by dividing its median fluorescence intensity (MFI) of 488B (red) by its MFI of 488C (green) and multiplying by 25% of the total resolution. The reported red/green ratio was then calculated by averaging each of the 100,000 cells’ ratios. Ratios are calculated from uncompensated linear data and are always reported as linear data. All ratios are only interpretable by comparison of the experimental condition with the positive and negative controls.

### Heterologous expression and purification of Cap1-C-terminal-His_6_-tagged construct

Full-length Cap1-C-terminal-His_6_-tagged construct was expressed in *E. coli* Rosetta^TM^ 2 (DE3) cells (Novagen). The freshly transformed cells were grown overnight as a primary culture at 37°C in Lysogeny broth (LB) media. The primary culture was further inoculated in terrific broth (TB) media and grown at 37°C till 1.2-1.3 OD_600nm_ and induced with 1 mM isopropyl β-D-1-thiogalactopyranoside (IPTG). The culture was further grown overnight at 16°C. The freshly harvested cells were resuspended in the lysis buffer (25 mM Hepes pH 8, 500 mM NaCl, 2 mM β-Mercaptoethanol and 5 % glycerol) supplemented with cOmplete mini, EDTA-free protease inhibitor tablets (Sigma). The resuspended cells were lysed by sonication at 50 % amplitude and 1 s on and 2 s off pulse for a total of 15 minutes. The unlysed cell debris were separated by centrifugation. The supernatant was used for harvesting the bacterial cell membrane. The membrane proteins were extracted from the purified bacterial cell membrane using 20 mM DDM (n-Dodecyl β-D Maltopyranoside) (Anatrace) detergent. The undissolved membrane was further separated using centrifugation and the supernatant was loaded on a pre-equilibrated (using lysis buffer with 1 mM DDM) 5 ml HisTrap column (Cytiva). The column was washed extensively with the lysis buffer supplemented with 40 mM imidazole and 1 mM DDM. The protein was eluted using the lysis buffer supplemented with 300 mM imidazole and 1 mM DDM. The protein fractions were checked on NuPAGE^TM^ 4-12 % Bis-Tris gel (Invitrogen) and the pure fractions were pulled together and concentrated to load on the Superdex200 10/300-increase column. The equilibration buffer and the running buffer used to perform size exclusion chromatography was 25 mM Hepes pH 8, 200 mM NaCl, 2 mM β-Mercaptoethanol, 5 % glycerol supplemented with 375 µM GDN (glycol-diosgenin, Anatrace) detergent.

### SEC-MALS analysis with Cap1-C-terminal-His_6_-tagged construct

The purified Cap1-C-terminal-His_6_-tagged sample was used for conjugate SEC-MALS analysis to determine the oligomeric state of the protein. The SEC-MALS experiment was performed using AKTA-Pure UV detector connected to SEC-MALS instrument (Wyatt) which has multi-angle light scattering detector and refractive index detector. Superdex200 10/300-increase column was used to run the protein sample in 25 mM Hepes pH 8, 200 mM NaCl, 2 mM β-Mercaptoethanol, 5 % glycerol and 375 µM GDN buffer. ASTRA 6 software was used for data analysis using protein conjugate method. The UV signal of the AKTA-Pure instrument was converted to analogue signal with a conversion factor of 1,000 mAU = 1 V. For protein conjugate analysis the refractive index increment (dn/dc) value for the protein was 0.185 ml g−1 and for the detergent was 0.14 ml g−1.

### Isothermal Titration Calorimetry (ITC) binding studies

ITC binding experiments were performed with 20 µM of His_6_-TEVsite-CARFL protein (the molarity was calculated with CARFL tetramer molecular weight) and its mutants titrated with 150 µM of cA_4_. The protein and the ligand were present in 25 mM Hepes pH 8, 200 mM NaCl, 2 mM β-mercaptoethanol and 5 % glycerol. MicroCal PEAQ-ITC (Malvern) instrument was used to perform the ITC studies at 25° C temperature. The protein was titrated with a total of 19 injections of cA_4_ out of which the volume of the first injection was 0.4 µl with a 0.8 s duration and rest of the 18 injections were 2 µl with a 4 s duration. The spacing between each of the injections was 150 s and the stirring speed was 750 rpm. The data was analyzed, and fitting was performed with MicroCal PEAQ-ITC Analysis software (Malvern) using one set of sites binding model. The estimated binding sites (N) value is 0.917 ± 1.4e-2. The estimated *K*_d_ value for the CARFL WT (wild type) was 35.6 ± 22.2 nM. The estimated ΔH (kcal/mol), ΔG (kcal/mol) and –TΔS (kcal/mol) values were –3.55 +/− 0.188, –10.2 and –6.61 respectively. The estimated *K*_d_ value for the CARFL (Q118A) mutant was 1.2 µM ± 0.6 µM. The estimated ΔH (kcal/mol), ΔG (kcal/mol) and –TΔS (kcal/mol) values were –5.07 ± 0.621, –8.07, –3 respectively. No binding curve could be fitted in the case of CARFL(Q119A) mutant. No binding was detected in the case of cA_3_ and cA_6_ ligands.

### Molecular dynamic (MD) simulations with cA4-bound Cap1

The initial system was built from the atomic coordinate file of cA_4_-bound Cap1 active state structure and parameterized using the CHARMM force field^50^, as provided by CHARMM-GUI. The system consisted of a protein complex of four protein chains, a single RNA molecule, and a lipid bilayer composed of 470 POPE and 152 POPG molecules. The system was solvated in a periodic tetragonal box with dimensions of 14.28 nm x 14.28 nm x 13.60 nm, containing 57,071 TIP3P^51^ water molecules. 150 mM KCl salt was added to the system. Molecular dynamics simulations were performed using GROMACS^52,53^. The system was energy-minimized using the steepest descent algorithm and equilibrated under canonical (NVT) and isothermal–isobaric (NPT) ensembles with gradually released positional restraints. Temperature was maintained at 303.15 K using the V-rescale thermostat, and pressure was controlled semi-isotropically at 1 bar using the Parrinello–Rahman barostat^54^. Production simulations were carried out for 100 ns with a 2 fs time step. Non-bonded interactions were computed using the Verlet cutoff scheme, with short-range electrostatic and van der Waals interactions truncated at 1.2 nm. Long-range electrostatics were treated using the Particle Mesh Ewald (PME)^55^ method and van der Waals interactions were smoothly switched off starting at 1.0 nm. All bonds involving hydrogen atoms were constrained using the LINCS algorithm^56^. Trajectories were saved every 100 ps for analysis.

To analyze selectivity mechanics, MD trajectories were evaluated across independent replicates (N = 3, 100 ns each) under KCl or NaCl conditions. The pore pathway was defined by residues E46, S49, Y52, D57, S61, and T64 and a 4.0 Å distance cutoff was used to analyze the incoming ions. *p* values were obtained with a two-sided paired t-test. Ion residence times within the Cap1 pore were quantified for K⁺, Na⁺, and Cl⁻ ions using trajectories from three independent molecular dynamics simulations conducted under KCl and NaCl conditions. Error bars indicate the mean ± s.e.m across three independent simulations. *p* values were obtained with two-sided t-tests with Welch’s correction.

### Cryo-EM sample preparation and imaging

The peak fraction of the purified apo-Cap1 protein was concentrated to 150 µM (calculated for monomeric Cap1) and supplemented with 800 µM of cA_4_ ligand (incubated for 30 minutes on ice). This sample was used for the cA_4_-Cap1 grid preparation. In the case of apo-Cap1, the protein was concentrated to 200 µM and used for grid preparation in the presence of 0.5 mM FOM (Fluorinated Octyl Maltoside) detergent. The UltratiFoil Au R (1.2/1.3) grids were used for both the samples. The grids were glow discharged for 2 minutes at 15 mA. Multiple grids were frozen at 4°C, 100 % humidity, 12 s wait time, 3.5 – 4.5 s blot time and 0 blot force using Vitrobot Mark IV (FEI). Both apo Cap1 and cA_4_-Cap1 datasets were collected using Krios G4 microscope at MSKCC equipped with a Falcon 4i detector and an energy filter of 10 eV slit width. The pixel size was 0.725 Å and the defocus range used for these datasets was –0.8 µm to –2.3 µm. The images were collected in the EER mode with a total electron dose of 59.33 electrons per Å^2^ with an exposure rate 11.5 electrons/pixel/sec. The EER unsampling factor was 1 and EER number of fractions were 45.

### Cryo-EM data processing and refinement

We used cryoSPARC v4.4.1^57^ for the processing of apo Cap1 and cA_4_-Cap1 datasets (Table S1).

#### Processing of apo-Cap1 tetramer dataset

We collected 11,929 movies and performed Patch Motion Correction and Patch CTF estimation jobs. 4,795,274 particles were picked using blob picker job with blob picking parameters 150 – 200 Å. The particles were extracted with a box size of 400 pixel.

Iterative 2D classification jobs were performed, and 207,755 particles were selected (Figs. S8A-B). Three ab-initio models were generated using ab-initio job (Fig. S8C). Out of the three models, the model (98,358 particles) marked by red box was used for the next refinement steps. Non-uniform refinement job was used to refine the map to 3.4 Å with C1 symmetry (Fig. S8D). The final set of particles used for the cryo-EM map was well distributed to different orientations (Fig. S8E) and the global resolution of the map was 3.4 Å following the standard FSC cutoff value of 0.143 (Fig. S8F). Further, local resolution values were estimated for the cryo-EM map of apo-Cap1 (Fig. S8G). The AF3^58^ models for each domain was used for the preliminary rigid body fitting using Chimera^59^ and the model was manually built using Coot^60^. Phenix^61^ real-space refinement program was used to remove the outliers and refine the model with a model vs. data correlation value (CC mask) of 0.84 (Table S1). In this cryo-EM map we observed densities to model two alternate conformations of the side chain of tyrosine 75 (Y75) residue of the TM1-TM2 segment (Figs. S9A-B). The model with flipped in Y75 side chain was refined with a model vs. data correlation value (CC mask) of 0.85 (Table S1). The model building for the cytosolic domain is displayed in Figures S6C-D.

#### Processing of cA_4_-Cap1 tetramer complex dataset

10,668 movies were collected for cA_4_-Cap1 sample and Patch Motion Correction and Patch CTF estimation jobs were performed using cryoSPARC. Blob picker job was used to pick up 42,443 particles from 1,061 micrographs. We selected 6,659 particles by 2D classification job to train a picking model using 1,061 micrographs to reduce the computation time using Topaz train job. The Topaz extract job was able to pick up 2,158,916 particles from 10,668 micrographs guided by previously trained model. 400 pixel box size was used for the particle extraction. Iterative 2D classification job was performed to select 325,786 particles (Figs. S10A-B). The particles were classified into three classes by 3D classification job and the class with 98,640 particles which showed all eight TM helices clearly was selected for the refinement job (Fig. 10C, red box). Non-uniform refinement was performed to resolve the map to 3.4 Å resolution with C1 symmetry (Fig. S10D). Another round of non-uniform refinement was performed with imposed C4 symmetry to improve the map quality and resolution specially at the TM domain (Fig. S10E). We observed good particle distribution both in the case of C1 and C4 symmetry maps (Figs. S10F-G). The standard FSC cut-off at 0.143 displayed 3.4 Å and 2.9 Å global resolution for C1 and C4 symmetry maps respectively (Figs. S10H-I). The cryo-EM map with C4 symmetry displayed that TM1-TM2 segment (Fig. S9E) and CARFL domain (Fig. S9F) was resolved with high resolution. Local resolution estimation on C1 symmetry map displayed that CARFL domain was resolved with high resolution even though the TM domain was poorly resolved (Fig. S10J). The local resolution estimation on the C4 symmetry-imposed map showed significantly improved resolution of the TM domain (Fig. S10K). AF3^58^ models of different domains were used for the rigid body fitting of the model using Chimera^59^. The model was further built using Coot^60^ and refined using real space refinement program in Phenix^61^ with a model vs. data correlation value (CC mask) of 0.85 (Table S1).

We re-processed 10,668 micrographs and picked up 4,629,563 particles using blob picker job in cryoSPARC. We extracted 3,838,293 particles using 400 pixel box size and performed iterative 2D classification jobs and screened 443,677 particles (Fig. S11A). Next, we built three ab-initio models out of which one model with 148,422 particles displayed the DUF domain (Figs. S11B-C). We used 3D classification job to classify 148,422 particles into five classes (Fig. S11D), out of which in one class the DUF domain was visible (Fig. S11D, red box). Next, we performed non-uniform refinement job with C1 symmetry which refined the map at 6.7 Å resolution (Fig. S11E). We imposed C4 symmetry in the next round of non-uniform refinement which improved the global resolution to 3.6 Å resolution (Fig. S11F). We observed good angular distribution for the particles used for final map generation (Fig. S11G). The FSC-plot showed 3.6 Å resolution at standard FSC (tight mask) cut-off 0.143 (Fig. S11H). The local resolution estimation of the map displayed even though the CARFL (Fig. S11G) and TM2 domain (Fig. S9H) was well resolved, the DUF has poor resolution (Fig. S11I). The TM1 and the loops were not resolved (Fig. S11I). We used AF3^58^ models for the rigid body fitting of the model using Chimera^59^. The model was further built using Coot^60^ and refined using real space refinement program in Phenix^61^ with a model vs. data correlation value (CC mask) of 0.73 (Table S1).

## Supporting information

Cap1 homologs

Strains, plasmids and oligonucleotides used in this study

## Acknowledgements

We thank Richard Hite and Yeeun (Jecy) Son for assistance during preliminary attempts at proteoliposome-based ion flux experiments. We thank Marianna Teplova for her efforts on ITC measurements. We thank Dr. Ahin Roy for his help in installation and standardizing the execution of the MD simulations on the Arvat computing facility at IIT Kharagpur. We thank M. J. de la Cruz and the MSKCC cryo-EM core facility staff for help with data acquisition. We thank S. Mazel, S. Semova, and S. Han from the Rockefeller Flow Cytometry Resource Center for their help in conceptualizing and running the flow cytometry experiment. LAM is supported by an NIH Award (R01 GM149834). LAM is an investigator of the Howard Hughes Medical Institute. DJP is supported by NIH (GM129430, AI141507 and GM145888), the Maloris Foundation and Memorial Sloan-Kettering Core grant (P30-CA008748). PM is supported by a Ludwig Cancer Research postdoctoral award (AWD00005337) and Faculty Start-up Research Grant (FSRG) of IIT, Kharagpur.

## Author contributions

PM, CWC, DJP and LAM conceived the study. CGR identified Cap1 and performed preliminary experiments. PM performed protein purification, cryo-EM, structural and molecular dynamics analysis, ITC experiments and all SEC-MALS analyses. CWC performed *in vivo* experiments. CFB performed sequence and structural homology searches and analyses. CWC and HC performed flow cytometry exeperiments for the membrane depolarization assay. PM and MT performed ITC experiments. PM, CWC, CFB, DJP and LAM wrote the manuscript. All authors read and approved the manuscript.

## Competing interests

LAM is a cofounder and Scientific Advisory Board member of Intellia Therapeutics, a cofounder of Eligo Biosciences and a Scientific Advisory Board member of Ancilia Biosciences.

## SUPPLEMENTARY FIGURE LEGENDS

**Figure S1.**
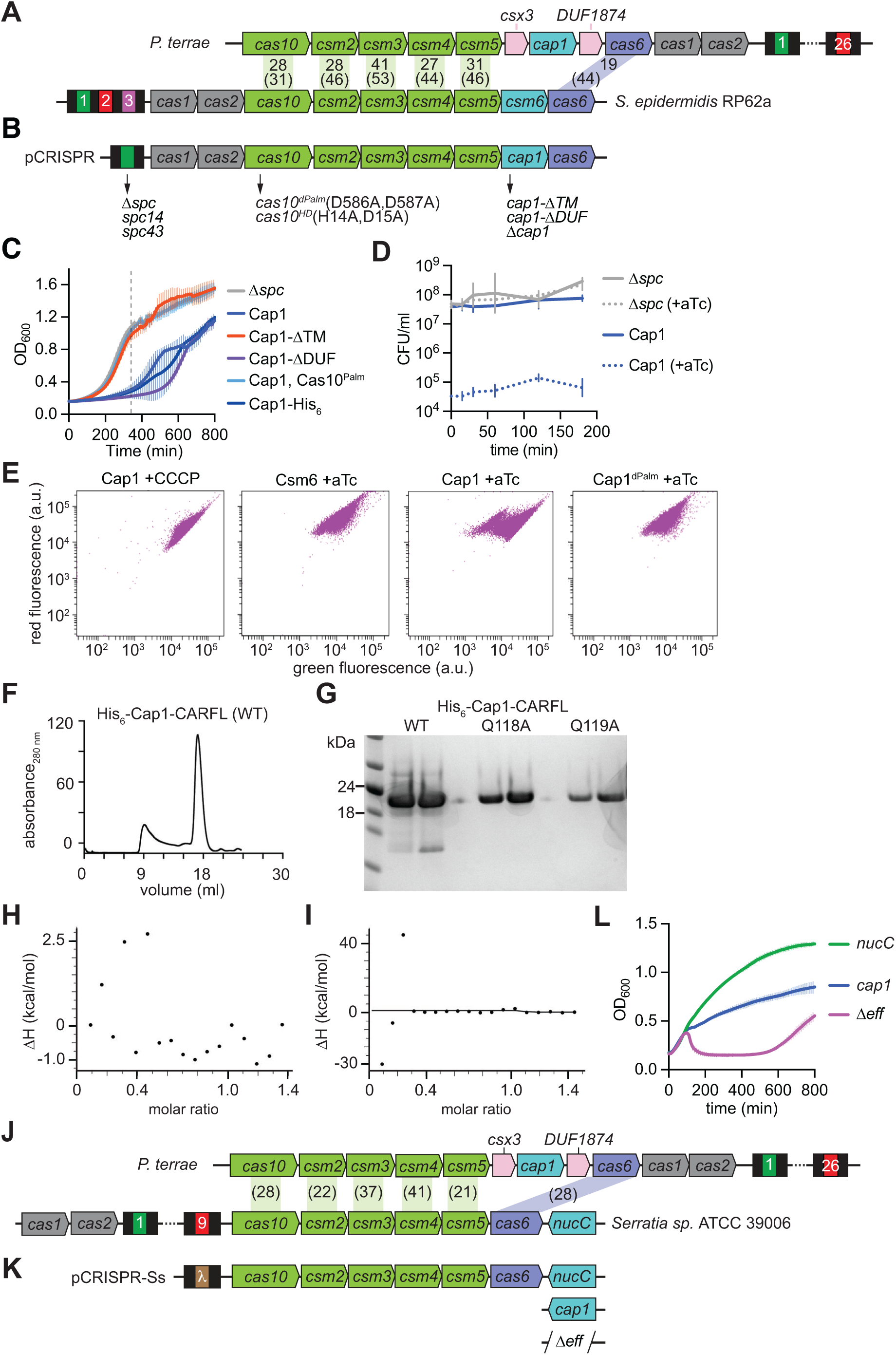
Cap1 activation and toxicity in staphylococci. **(A)** Comparison of the III-A systems of *P. terrae* and *S. epidermidis* RP62a. Black boxes indicate CRISPR repeats; colored, numbered boxes indicate CRISPR spacers. Numbers indicate the percent homology or identity (in parentheses) of conserved Cas proteins. **(B)** Genetic modifications of the *S. epidermidis* RP62a type III-A CRISPR locus cloned into various pCRISPR plasmids. Amino acid substitutions, domain deletions, and insertion of different spacer sequences are indicated. **(C)** Growth of staphylococci carrying pTarget and diverse pCRISPR variants, measured as OD_600_ after the addition of aTc. Dotted line crosses at 360 minutes, the time used to generate bar graphs that represent the the different growth rates. The mean of three biological replicates +/− s.e.m are shown. **(D)** Enumeration of CFU within staphylococcal cultures carrying pTarget and pCRISPR(Cap1) or the control version of this plasmid without a targeting spacer (Δ*spc*). Aliquots taken at different times after addition of aTc to induce target transcription and cA_4_ synthesis were plated on plasmid-selective media with or without aTc. **(E)** Flow cytometry of *S. aureus* cells harboring various pCRISPR constructs and stained with DiOC2(3), collected 30 minutes after addition of aTc or the depolarizing agent CCCP. “a.u.”, arbitrary units; green fluorescence, emission of 515 nm upon 488C excitation; red fluorescence, emission of 586 nm upon 488B excitation. Each plot is representative of approximately 100,000 cells. **(F)** Chromatogram obtained after size exclusion purification of wild-type His_6_-Cap1-CARFL. **(G)** SDS-PAGE followed by Coomassie-blue staining of purified wild-type and mutant His_6_-Cap1-CARFL proteins. **(H)** ITC experiment of purified His_6_-Cap1-CARFL performed with cA_3_. **(I)** Same as (H) but for cA_6_. **(J)** Same as (A) but comparing the type III-A CRISPR-Cas systems of *P. terrae* and *Serratia sp*. ATCC 39006. **(K)** Same as (B) but for the cloning of the type III-A CRISPR-Cas system of *Serratia sp.* ATCC 39006 to generate pCRISPR-Ss plasmids. “λ” indicates a spacer that matches a sequence within the gene *O* of lambda phage. **(L)** Growth of *E. coli* carrying different pCRISPR-Ss constructs, measured as OD_600_ after infection with lambda phage at MOI ∼0.5. The mean of three biological replicates +/− s.e.m are shown.

**Figure S2.**
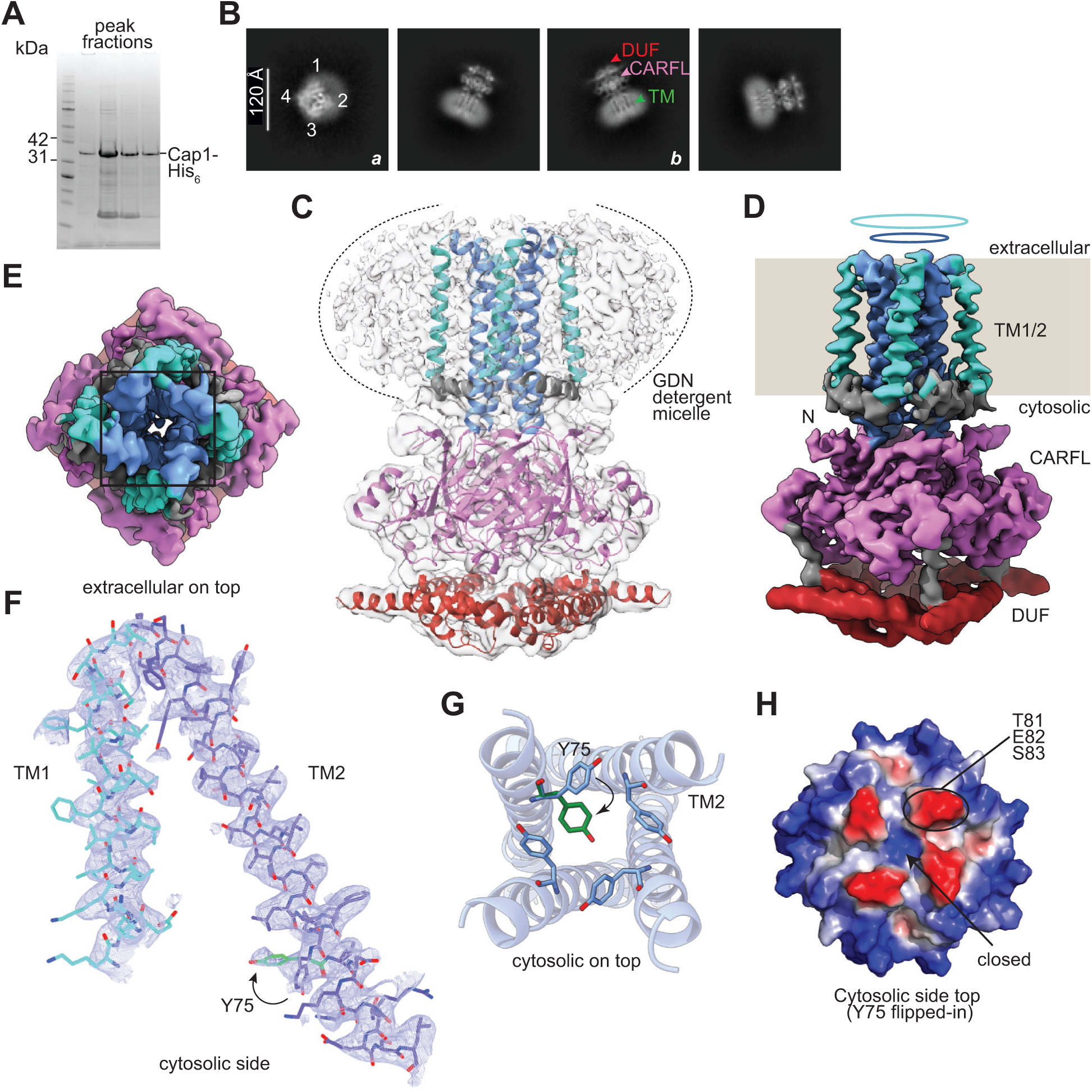
Apo-Cap1 structure. **(A)** SDS-PAGE followed by Coomassie-blue staining of peak fractions of purified full-length apo-Cap1 (34 kDa). **(B)** Representative 2D class averages showing different views of the apo-Cap1 protein. Class *a* displays a front view with cytosolic domains on top. Class *b* displays a side view in which the DUF4579, CARFL and TM1/2 domains are labelled. **(C)** Ribbon diagram of the apo-Cap1 tetramer showing in light grey the density for the GDN detergent micelle surrounding the transmembrane domains; the detergent density is marked by dotted lines. **(D)** Cryo-EM map of the apo-Cap1 tetramer. TM1 and TM2 are displayed in green and blue and form outer and inner rings, respectively, that insert into the bacterial membrane (beige background, which replaces the density observed for the GDN detergent micelle). The N-terminal segment (N, grey), CARFL domain (pink), the interconnecting loop (L, grey) and the DUF4579 domain (red) locate on the cytosolic side. **(E)** Cryo-EM map of apo-Cap1 viewed from the extracellular side showing extra density inside the pore, protruding inwards from one of the inner TM2 helices. The inner TM2 ring is delimited by the black box. **(F)** Density for the TM1 and TM2 helices shown in blue and green mesh; the black arrow indicates the two alternate conformations of the Y75 side chain. **(G)** Tetrameric arrangement of the inner TM2 ring viewed from the cytosolic side displaying the two alternate conformations of Y75 for one out of the four TM2 helices. The flipped Y75 side chain is shown in green. **(H)** Electrostatic surface representation of the cytosolic side view of the TM1 and TM2 (with Y75 side chain flipped in) domains, showing a closed pore.

**Figure S3.**
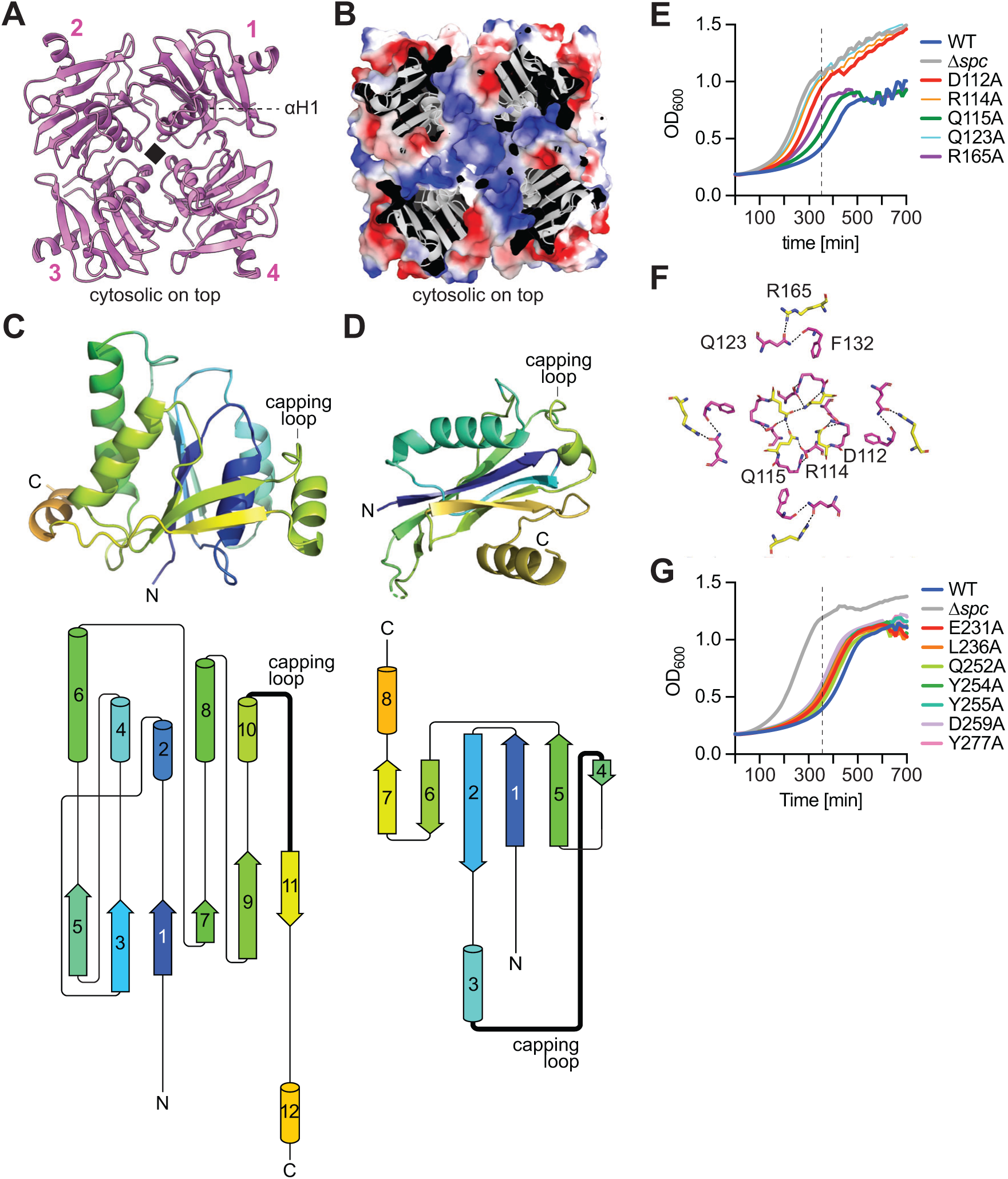
Unique properties of Cap1’s CARFL domain. **(A)** Cytosolic view of the tetrameric CARFL domain of apo-Cap1, arranged in 4-fold symmetry. Alpha helix 1 (αH1) of one of the CARFL domain is labelled. **(B)** Electrostatic surface representation of the cytosolic side view of the tetrameric CARFL of apo-Cap1, showing a positively charged pocket at the center. **(C)** Top; conventional CARF monomer domain from apo-Cad1 protein (PDBID 9C77) showing a β-sheet formed by five parallel β-strands (1, 3, 5, 7, 9) and one antiparallel β-strand (11) sandwiched between of six alpha helices. Bottom; 2D topology map of the top structure. **(D)** Top; CARFL monomeric domain of apo-Cap1 showing a β-sheet composed of three parallel β-strands (1, 5, 7) and three antiparallel β-strands (2, 4, 6) sandwiched between a pair of alpha helices, 3 and 8. Bottom; 2D topology map of the top structure. **(E)** Growth of staphylococci carrying pTarget and diverse pCRISPR variants harboring alanine substitutions of Cap1 residues shown in Figure 2F, measured as OD_600_ after the addition of aTc. Dotted line crosses at 360 minutes, the time used to generate bar graphs that represent the the different growth rates. The mean of three biological replicates +/− s.e.m are shown. **(F)** Interacting residues of the apo-CARFL tetramer showing interactions between the side chains of residues Q123, R165, F132, D112, R114 and Q115. **(G)** Growth of staphylococci carrying pTarget and diverse pCRISPR variants harboring alanine substitutions of Cap1 residues shown in Figure 2I, measured as OD_600_ after the addition of aTc. Dotted line crosses at 360 minutes, the time used to generate bar graphs that represent the different growth rates. The mean of three biological replicates +/− s.e.m are shown.

**Figure S4.**
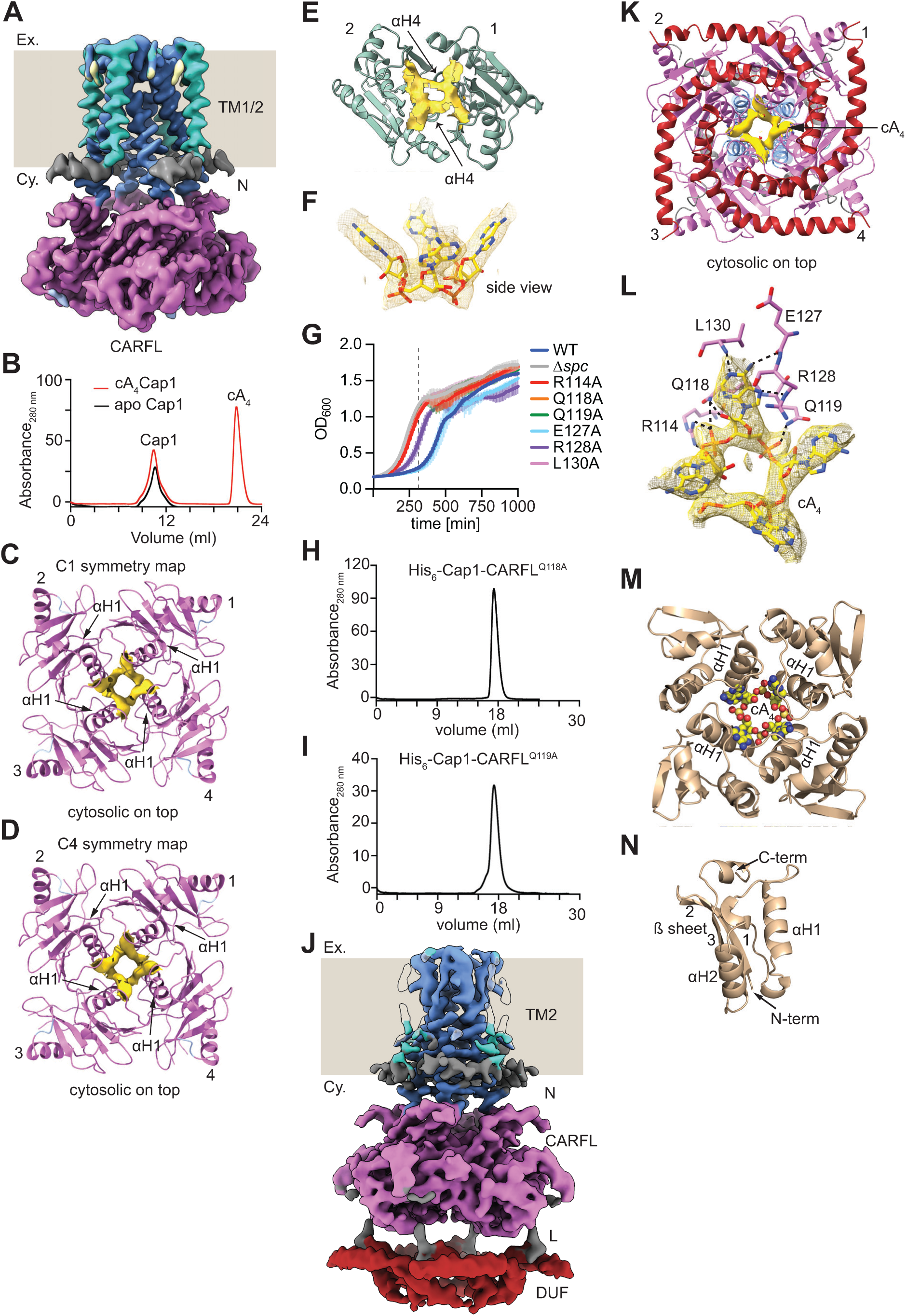
cA_4_-bound Cap1 structure. **(A)** Cryo-EM map of the cA_4_-Cap1 tetramer. TM1 and TM2 are displayed in green and blue and form outer and inner rings, respectively, that insert into the bacterial membrane (beige background, which replaces the density observed for the GDN detergent micelle; “Ex.”, extracellular space, “Cy.”, cytosol). The N-terminal segment (N, grey) and the CARFL domain (pink) locate on the cytosolic side. Density for the DUF4579 domain was not detected. **(B)** Size exclusion chromatograms of purified cA_4_-bound Cap1 (red) and apo-Cap1 (black). The peak corresponding to unbound excess cA_4_ is labelled. **(C)** Structure of the tetrameric CARFL domain of cA_4_-bound Cap1 (without visible DUF4579) displaying the αH1 helices forming the cA_4_ binding pocket. The density of the cA_4_ molecule is shown in yellow from C1 symmetry map. **(D)** Same as (C) with the cA_4_ density displayed from a C4 symmetry-imposed map. **(E)** Cad1-CARF dimer with bound cA_4_ (PDB ID: 9C77). The base of the cA_4_ binding pocket is formed by a pair of αH4 helices of each monomer, indicated by black arrows. **(F)** Side view of bound cA_4_ within the CARFL pocket, showing adenine bases pointing upwards, on a different plane than the rest of the molecule. **(G)** Growth of staphylococci carrying pTarget and diverse pCRISPR variants harboring alanine substitutions of Cap1 residues shown in Figure 3C, measured as OD_600_ after the addition of aTc. Dotted line crosses at 360 minutes, the time used to generate bar graphs that represent the different growth rates. The mean of three biological replicates +/− s.e.m are shown. **(H)** Chromatogram obtained after size exclusion purification of Cap1-CARFL^Q118A^-His_6_. **(I)** Same as **(H)** for Cap1-CARFL^Q119A^-His_6_. **(J)** Cryo-EM map of the cA_4_-Cap1 tetramer. TM1 is not visible. TM2 is displayed in blue and forms an inner ring that inserts into the bacterial membrane (beige background, which replaces the density observed for the GDN detergent micelle; “Ex.”, extracellular space, “Cy.” cytosol). The N-terminal segment (N, grey), CARFL domain (pink), the linker (L, grey) and the DUF4579 domain (red) locate on the cytosolic side. **(K)** Cytosolic view of the cA_4_-bound Cap1 structure with a visible DUF4579. cA_4_ (yellow) can be seen bound to the center of 4-fold symmetry of the CARFL domain, behind the DUF4579 helix-turn-helices tetramer. **(L)** CARFL domain residues interacting with cA_4_ in the cA_4_-bound Cap1 structure with a visible DUF4579. **(M)** Tetrameric arrangement of C-terminal cA_4_ binding domain of Csx23 (PDB 8QJK) with bound ligand. **(N)** The domain architecture of Csx23 protomer.

**Figure S5.**
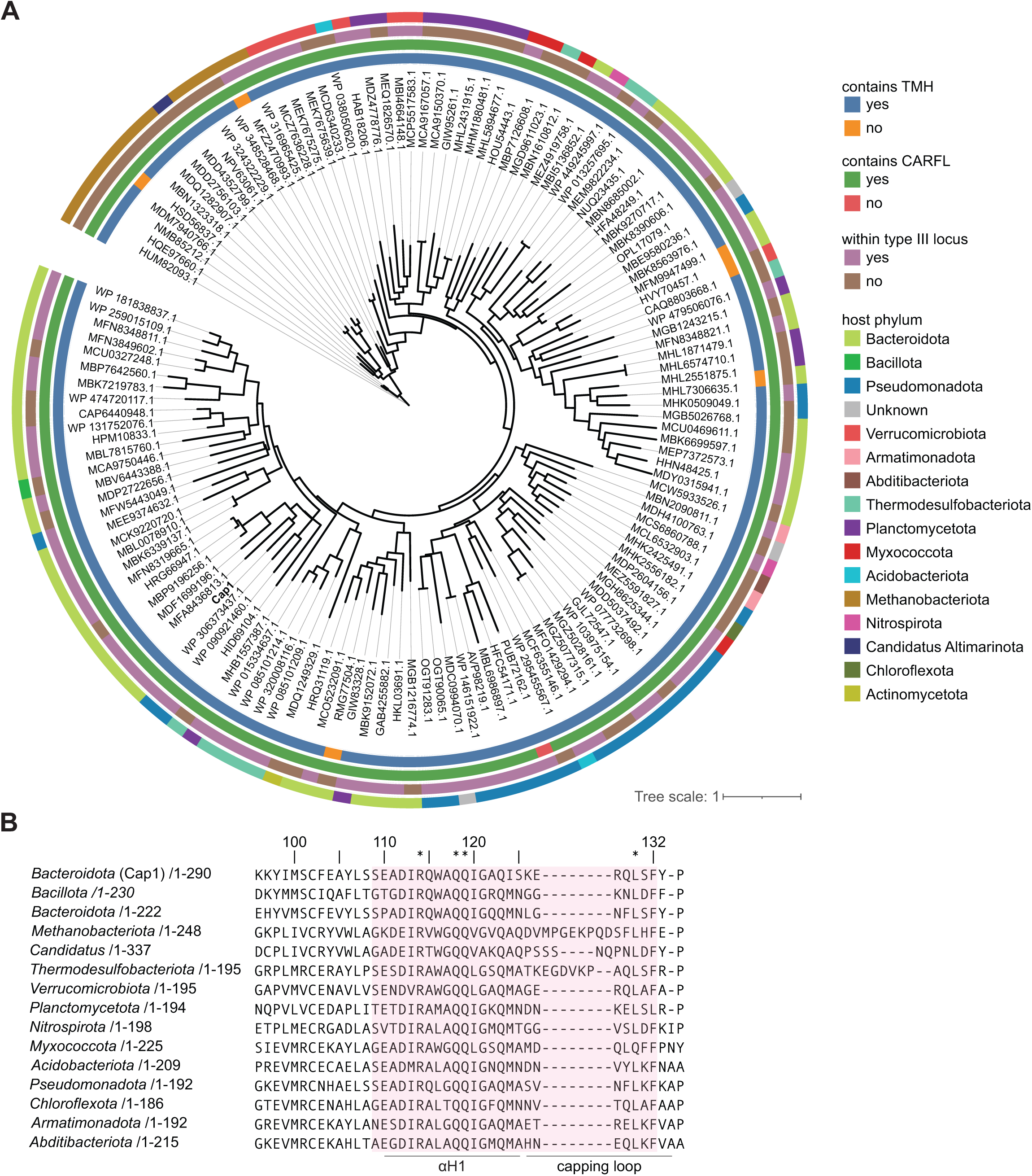
PtCap1 sequence homologs. **(A)** Phylogenetic tree built with PtCap1 protein sequence (accession number WP 090655733.1) homologs. Four properties of each homolog are indicated: the presence of a TMH domain, the presence of CARFL, whether it was found within a type III CRISPR-Cas locus, and the host phylum. **(B)** Multiple sequence alignment of CARFL domains from Cap1 homologs across diverse bacterial phyla. The residues from the αH1 and capping loop structural elements are highlighted in red. The conserved residues within the cA_4_-binding motif, R(X_3_)QQ(X_10_)L, are noted with an asterix.

**Figure S6.**
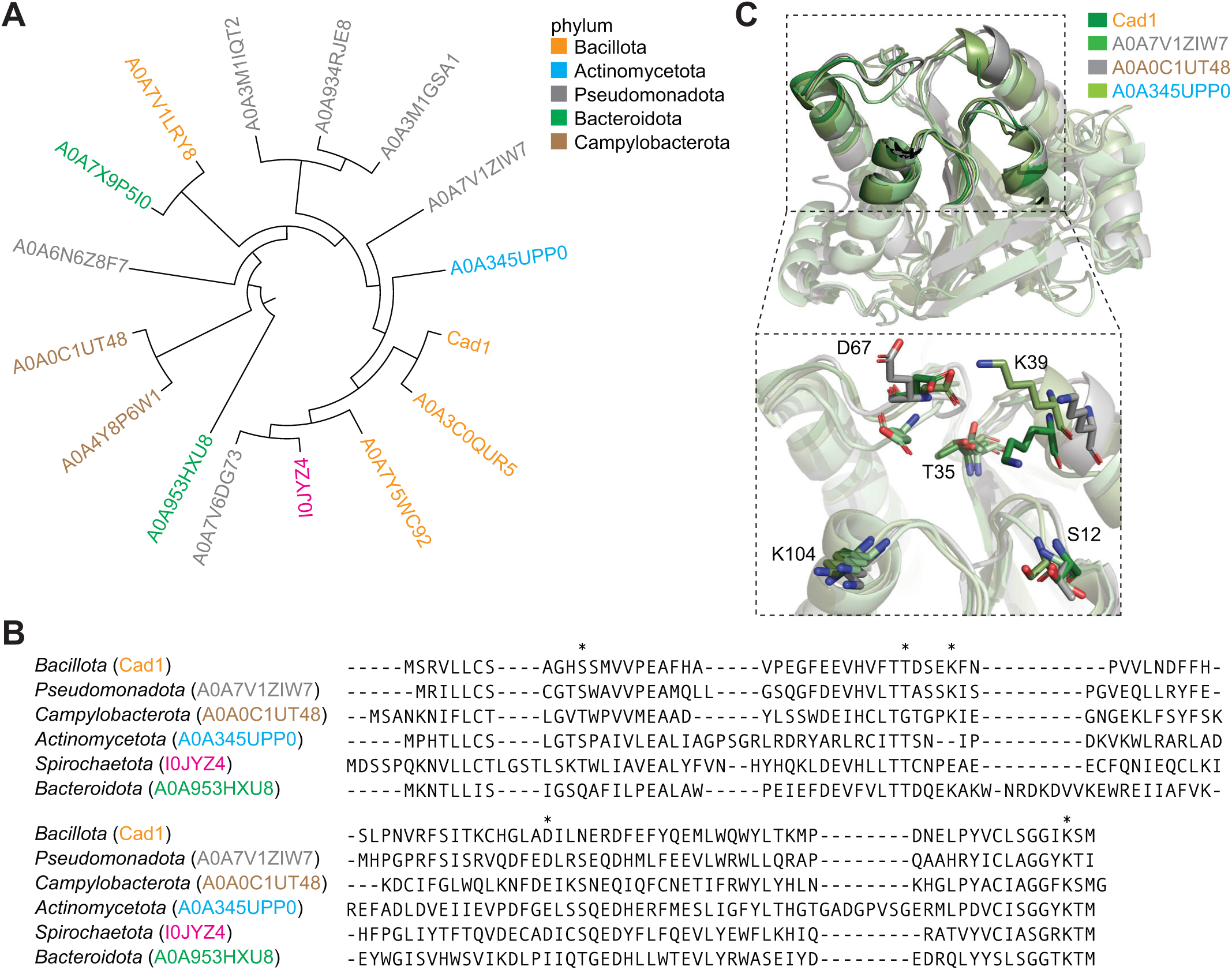
Cad1 structural homologs. **(A)** Phylogenetic tree based on structural homology search using Cad1 (PDB 9C77) CARF. The top 15 hits (E value 2.5e-17 to 8.20e-10) from the AFDB50 database are presented and colored according to the phyla of their host. **(B)** Multiple sequence alignment of one representative sequence from each bacterial phyla is presented. The cA_4_/cA_6_ binding residues are marked by an asterix. **(C)** Structural alignment of Cad1 CARF and three of its structural homologs. The inset shows a detailed view of the cA_4_/cA_6_ binding residues across the different homologs.

**Figure S7.**
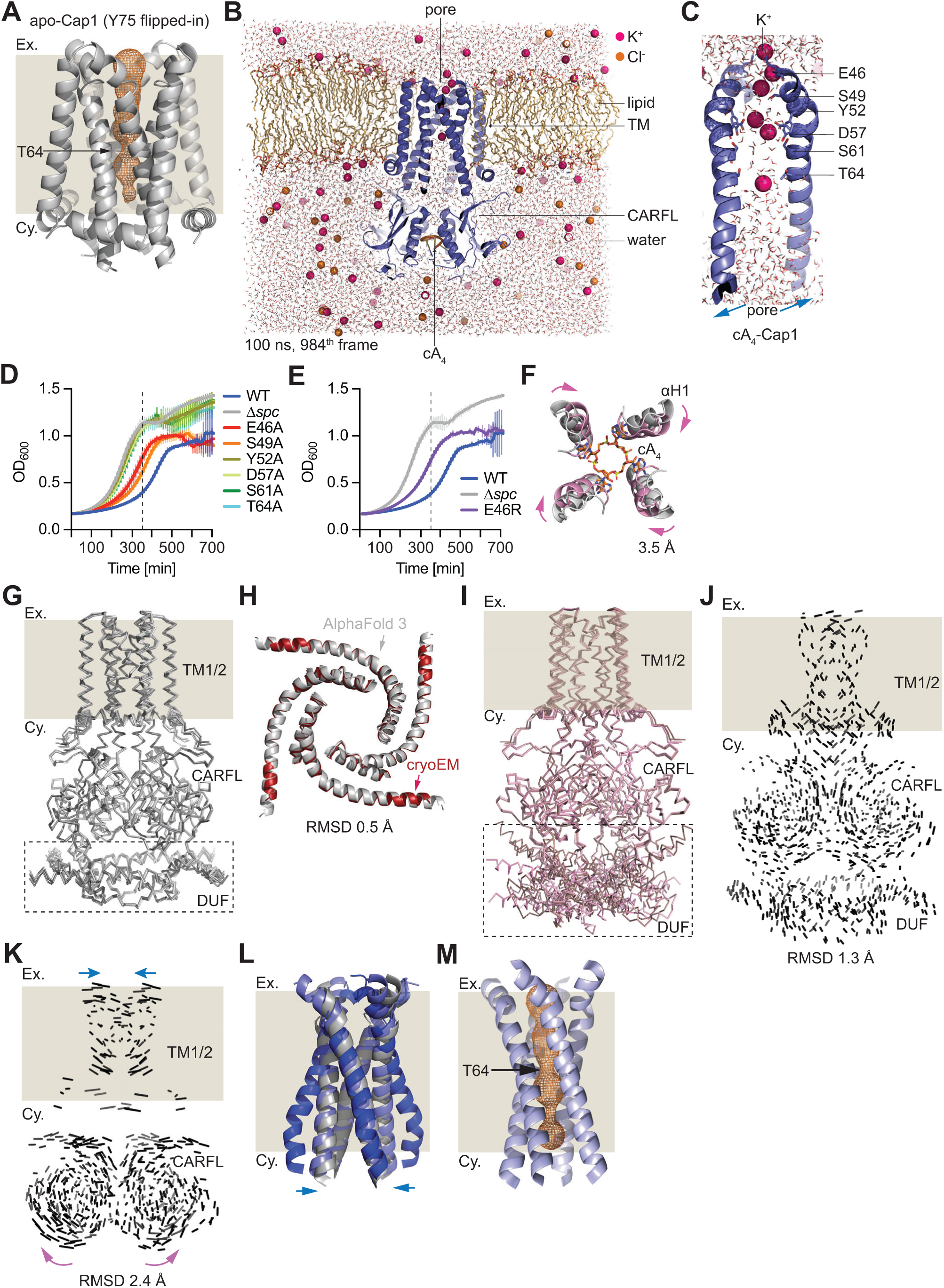
Conformational changes of Cap1 upon cA_4_ binding lead to pore opening and membrane depolarization. **(A)** Pore within the transmembrane region of apo-Cap1 structure (Y75 side chain flipped-in), predicted using Mole2.5 software. The bacterial membrane is represented by the beige background; “Ex.”, extracellular space, “Cy.”, cytosol. The inner volume of the pore is displayed by the orange mesh. The narrowest part of the pore is surrounded by four T64 residues belonging to inner TM2 helices. The predicted pore fails to cross the full membrane layer. **(B)** MD simulation (100 ns) trajectory at 984^th^ frame showing the distribution of potassium and and absence of chloride ions across the membrane in the presence of cA_4_-bound Cap1. **(C)** Enlarged view of the pore from **(B)**, highlighting potassium ions in close proximity to negatively charged and polar residues lining the inner TM2 helix. **(D)** Growth of staphylococci carrying pTarget and diverse pCRISPR variants harboring alanine substitutions of Cap1 residues shown in Figure 4G, measured as OD_600_ after the addition of aTc. Dotted line crosses at 360 minutes, the time used to generate bar graphs that represent the the different growth rates. The mean of three biological replicates +/− s.e.m are shown. **(E)** Same as **(D)** for the E46R substitution. The mean of three biological replicates +/− s.e.m are shown. **(F)** Inward movement of αH1 helices of the CARFL domain upon cA_4_ binding, indicated by magenta arrows. **(G)** Superposition of the top four AF3 models for apo-Cap1, showing a good agreement across all domains, including the DUF4579 domain (dotted rectangle). The bacterial membrane is represented by the beige background; “Ex.”, extracellular space, “Cy.”, cytosol. **(H)** Superposition of the helix-turn-helices of the tetrameric DUF4579 experimentally determined by cryo-EM (red) of apo-Cap1 with the structure predicted by the top AF3 model (grey), showing a good alignment with RMSD 0.5 Å. **(I)** Superposition of the top four AF3 models for cA_4_-Cap1, showing a good agreement across all domains, except the DUF4579 domain that displayed heterogenous structures (dotted rectangle). The bacterial membrane is represented by the beige background; “Ex.”, extracellular space, “Cy.”, cytosol. **(J)** Superposition of cA_4_-Cap1 structure with visible DUF4579 and apo-Cap1, showing a 1.3 Å RMSD value. The bacterial membrane is represented by the beige background; “Ex.”, extracellular space, “Cy.”, cytosol. **(K)** Superposition of cA_4_-Cap1 with and without visible DUF4579, showing a 2.4 Å RMSD value. The bacterial membrane is represented by the beige background; “Ex.”, extracellular space, “Cy.” Cytosol. Blue arrows, inward movement of TM1/2 domain; pink arrows, outward movement of the CARFL domain. Arrows show movement of cA_4_-Cap1 structure with a visible DUF4579, relative to cA_4_-Cap1 lacking structured DUF4579. **(L)** Comparison of TM2 domains of apo (grey), cA_4_-Cap1 with visible DUF4579 (light blue) and cA_4_-Cap1 without visible DUF4579 (dark blue). The bacterial membrane is represented by the beige background; “Ex.”, extracellular space, “Cy.”, cytosol. Blue arrows show TM2 helices moving inward in DUF4579–cA_4_-Cap1 relative to cA_4_-Cap1 lacking structured DUF4579. **(M)** Same as (A) for the cA_4_-Cap1 with visible DUF4579 structure.

**Figure S8.**
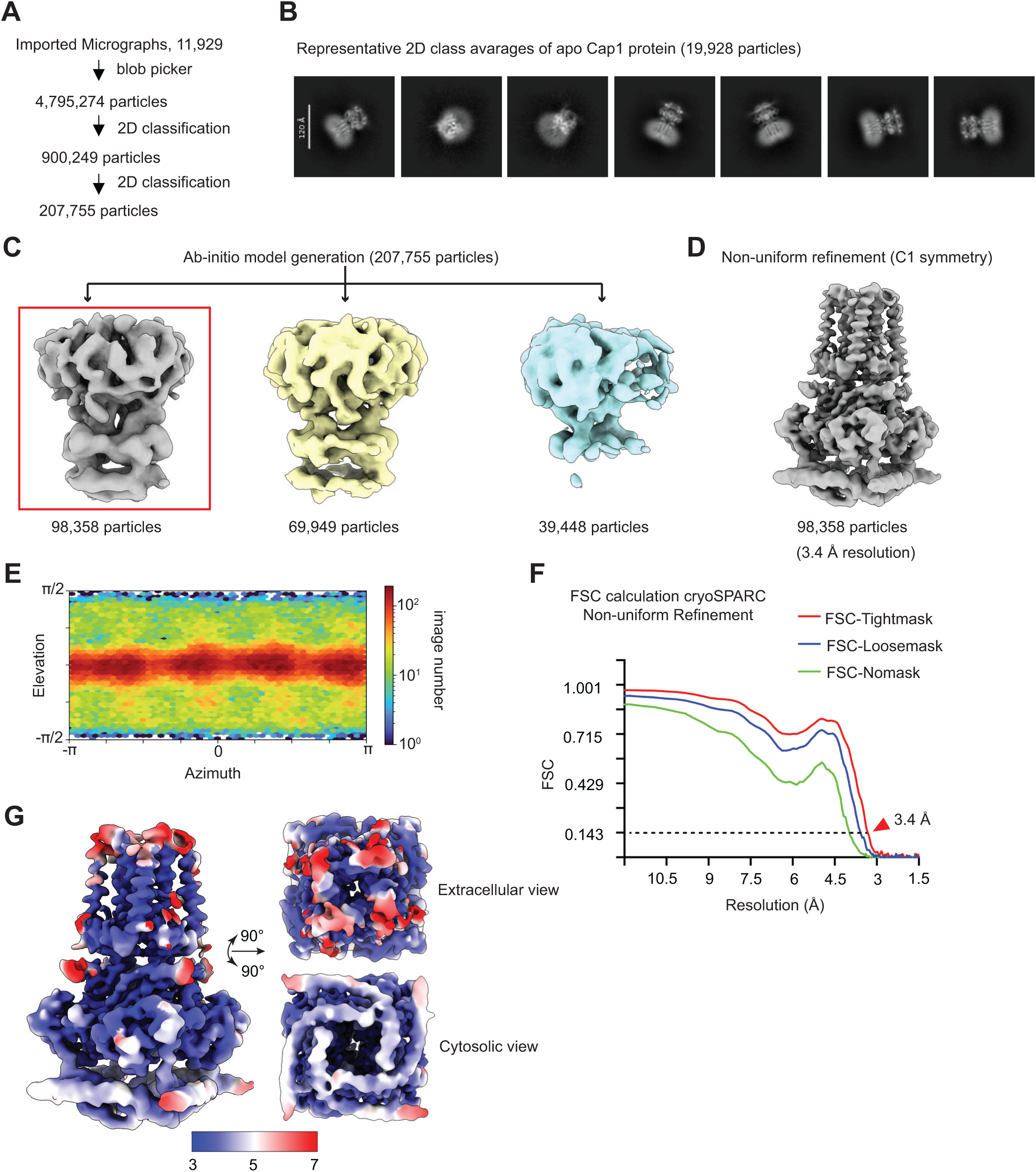
Cryo-EM data processing for apo-Cap1 protein. **(A)** Data processing steps including particle picking and 2D classification for the apo-Cap1 protein sample. The particle numbers are mentioned for each step. **(B)** Representative 2D class averages with different views of the apo-Cap1 protein. **(C)** Three ab-initio models were generated with final selected particles by 2D classification. The first model was selected for next round of refinement process (red box). **(D)** Cryo-EM map refined by non-uniform refinement job with C1 symmetry. **(E)** Angular distribution of the particles used for the final map. **(F)** Fourier shell correlation (FSC) plot, with and without mask. FSC value of 0.143 was used as threshold for resolution determination. **(G)** Local resolution estimated map displayed in different views including side view, extracellular view and cytosolic view. The scale bar is represented in Å unit.

**Figure S9.**
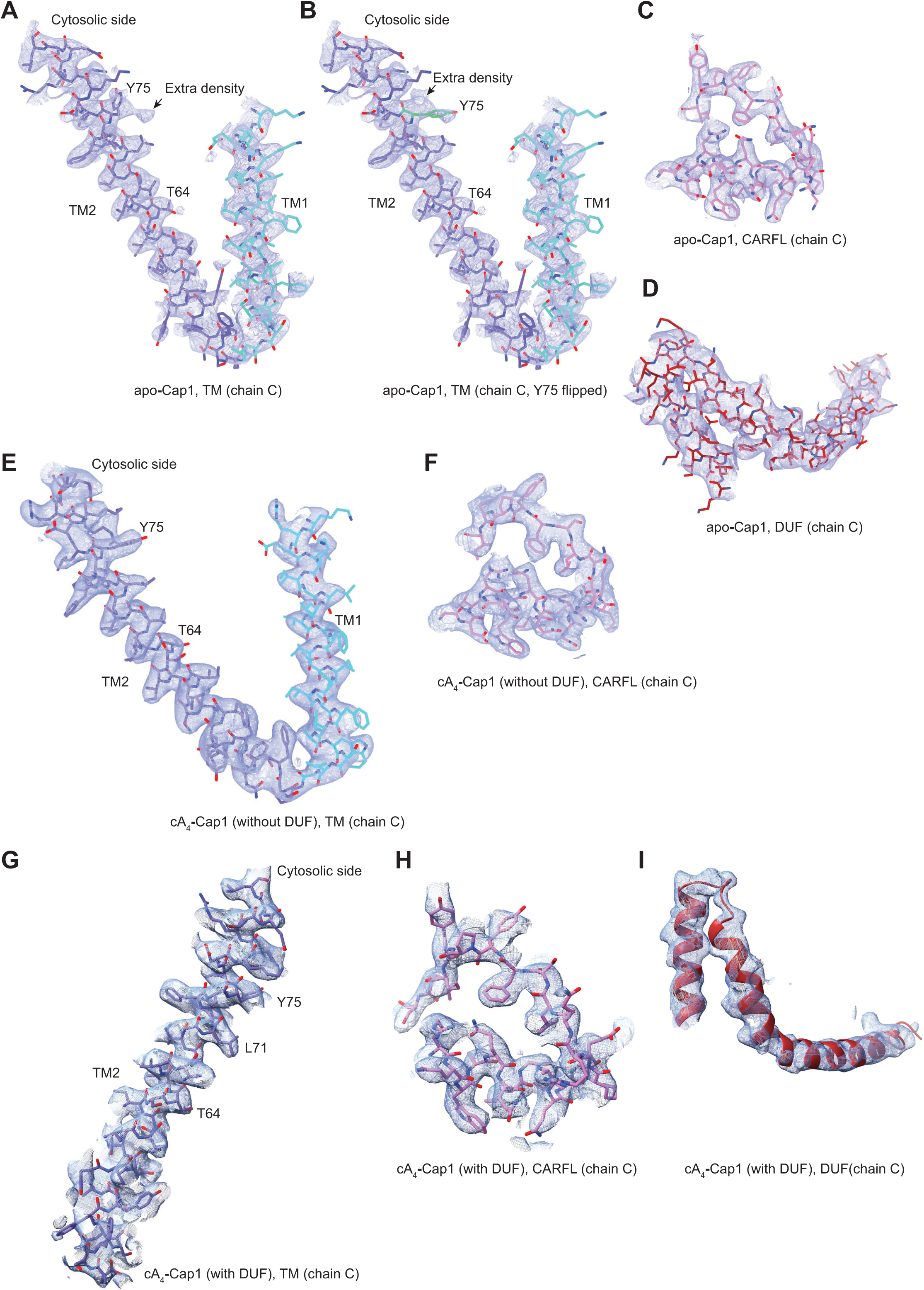
Density tracing in the cryo-EM map of apo– and cA_4_-bound Cap1 protein. **(A)** Map of the apo-Cap1 TM1 and TM2 segments indicating the extra density for the alternate conformation of Y75 side chain (black arrow) at contour level ∼5 RMSD. T64 residue is labelled. **(B)** Same density map as (A) displaying Y75 side chain at flipped-in conformation. The flipped side chain of Y75 is colored in green. **(C)** Electron density map of part of the apo-Cap1 CARFL domain displayed at contour level ∼5 RMSD. **(D)** Same as (C) displaying for the apo Cap1 DUF4579 domain. **(E)** Electron density map of cA_4_-Cap1 (without visible DUF4579) C4 symmetry-imposed map at contour level ∼5 RMSD displaying representative TM1 and TM2 segments. Y75 and T64 residues are labelled. **(F)** Similar representation as in panel (E) showing map for part of the CARFL domain at contour level ∼5 RMSD. **(G)** Map of one of the TM2 domain from cA_4_-Cap1-DUF4579 structure displayed at contour level ∼5 RMSD. **(H-I)** Similar representation of the maps as panel (G) for the CARFL and DUF at similar contour level.

**Figure S10.**
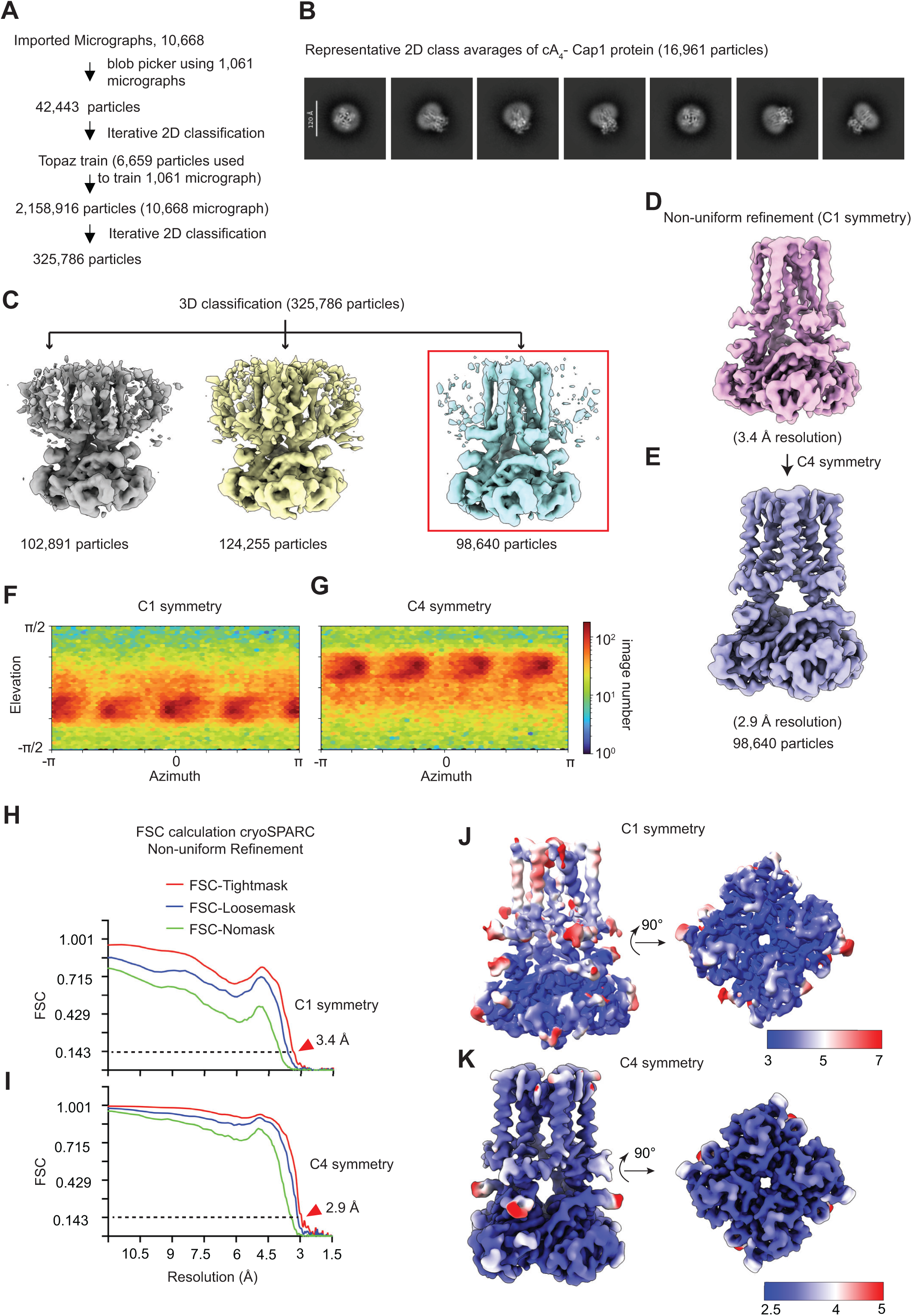
Cryo-EM data processing for cA_4_-Cap1 (without visible DUF) complex. **(A)** Steps used for cA_4_-Cap1 data processing including particle picking by blob picker and Topaz, and 2D classification jobs. **(B)** Representative 2D class averages are displayed with different views of cA_4_-Cap1 complex. **(C)** Cryo-EM maps generated using 3D classification job. The class marked with red box was used for next round of refinement. **(D)** Cryo-EM map of the cA_4_-Cap1 complex refined by non-uniform refinement job with C1 symmetry. **(E)** The resolution of the map shown in (D) was further improved by another round of non-uniform refinement with C4 symmetry. **(F)** Angular distribution of the particles used in the non-uniform refinement with C1 symmetry. **(G)** Similar plot as in (F) in the case of the non-uniform refinement with C4 symmetry in displayed. **(H)** FSC plot with and without mask generated by non-uniform refinement with C1 symmetry. **(I)** Similar FSC plot as in (H) generated by non-uniform refinement with C4 symmetry. **(J)** Local resolution estimation of the C1 symmetry cryo-EM map of cA_4_-Cap1 complex. **(K)** Similar as in (J) for C4 symmetry cryo-EM map of the cA_4_-Cap1 complex. The scale bar is presented in Å unit for both maps in (J) and (K).

**Figure S11.**
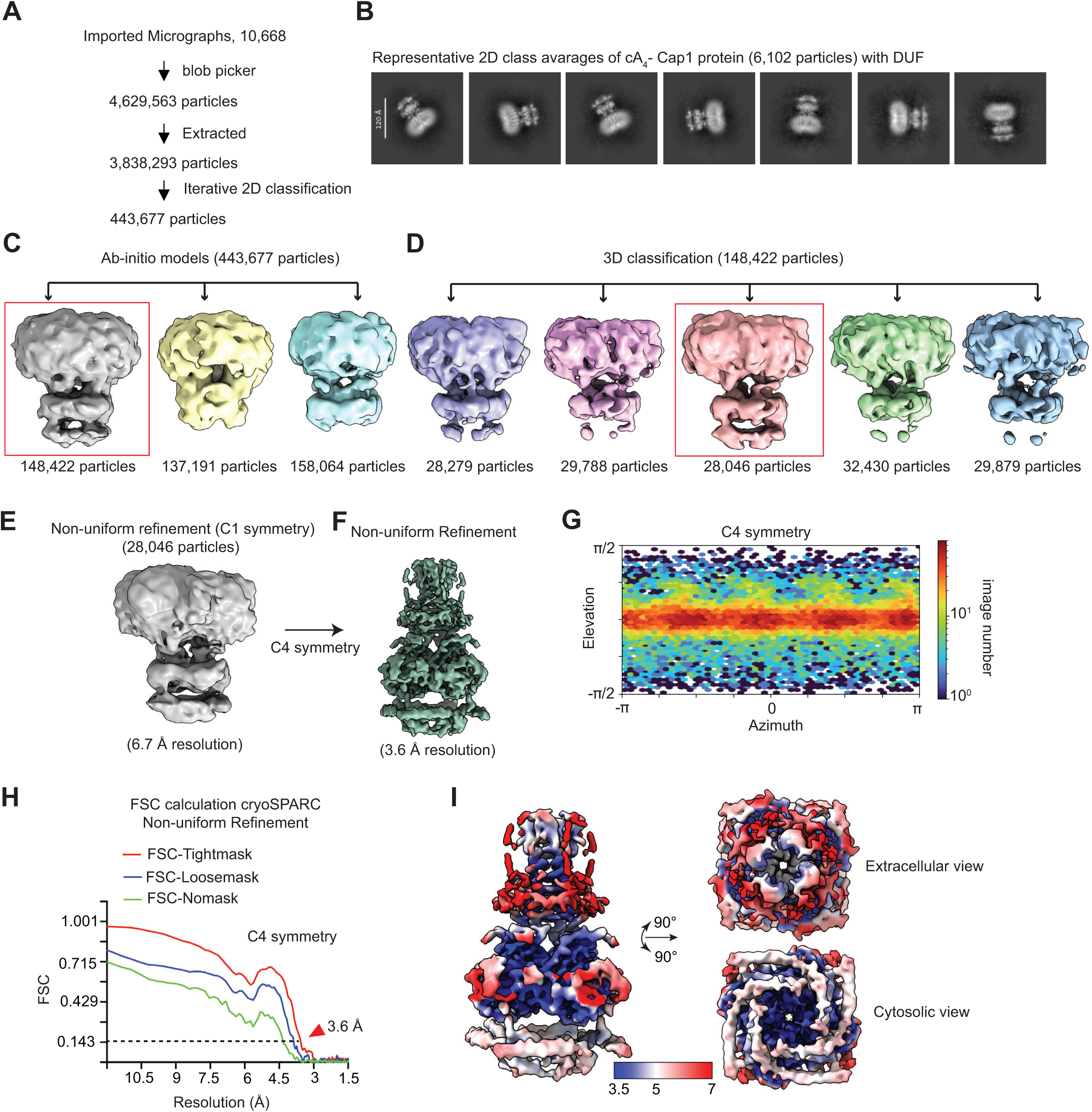
Cryo-EM data processing for cA_4_-Cap1 (with visible DUF4579) complex. **(A)** Particles were picked up using blob picker and screened using iterative 2D classification job. **(B)** Representative 2D classes (6,102 particles) are displayed with visible DUF4579 domain. **(C)** The particles selected by 2D classification job were used to build ab-initio models. A model marked by red box displayed the DUF4579 domain. **(D)** This model corresponding to 148,422 particles was used for 3D classification job. Out of the five 3D classes, one class containing 28,046 particles displayed intact DUF4579 domain marked in red box. **(E)** Non-uniform refinement job was performed with C1 symmetry, that resolved the map at 6.7 Å resolution. **(F)** C4 symmetry imposed non-uniform refinement improved the global resolution to 3.6 Å. **(G)** Angular distribution of the particles used for the refinement jobs. **(H)** FSC plot generated by the non-uniform refinement job. **(I)** The local resolution estimation indicates that the CARFL and TM2 domains have resolution close to 3.5 Å but the DUF4579 domain was poorly resolved. Additionally, TM1 and the loops were not resolved. The scale bar is presented in Å unit.

**Table S1.** Cryo-EM data collection, refinement, and validation statistics.

|  | Apo-Cap1<br>(Y75 flipped<br>out)<br>(PDB 9Z6G)<br>(EMDB-73839) | Apo-Cap1<br>(Y75 flipped<br>in)<br>(PDB 9Z6H)<br>(EMDB-73839) | cA<sub>4</sub>-Cap1<br>(PDB 9Z6J)<br>(EMDB-73841) | cA<sub>4</sub>-Cap1 with<br>DUF<br>(PDB 9Z6K)<br>(EMDB-73842) |
| --- | --- | --- | --- | --- |
| **Data collection and Processing** |  |  |  |  |
| Microscope | Krios G4 | Krios G4 | Krios G4 | Krios G4 |
| Voltage (keV) | 300 | 300 | 300 | 300 |
| Camera | Falcon 4i | Falcon 4i | Falcon 4i | Falcon 4i |
| Magnification | 165000 | 165000 | 165000 | 165000 |
| Pixel size at detector (Å/pixel) | 0.725 | 0.725 | 0.725 | 0.725 |
| Total electron exposure (e<sup>-</sup>/Å<sup>2</sup>) | 59.33 | 59.33 | 59.33 | 59.33 |
| Exposure rate (e-/pixel/sec) | 11.5 | 11.5 | 11.5 | 11.5 |
| EER number of fractions | 45 | 45 | 45 | 45 |
| EER unsampling factor | 1 | 1 | 1 | 1 |
| Defocus range (µm) | -0.8 to -2.3 | -0.8 to -2.3 | -0.8 to -2.3 | -0.8 to -2.3 |
| Phase plate (if used) | - | - | - | - |
| - phase shift range (in degrees) | - | - | - | - |
| - number of images per phase plate | - | - | - | - |
| Automation software | EPU | EPU | EPU | EPU |
| Tilt angle (if grid was tilted) | - | - | - | - |
| Energy filter slit width (eV) | 10 | 10 | 10 | 10 |
| Micrographs collected (no.) | 11,929 | 11,929 | 10,668 | 10,668 |
| Micrographs used (no.) | 11,929 | 11,929 | 10,668 | 10,668 |
| Total extracted particles (no.) | 4,795,274 | 4,795,274 | 2,158,916 | 4,629,563 |
| <u>For each reconstruction:</u> |  |  |  |  |
| Refined particles (no.) | 98,358 | 98,358 | 98,640 | 28,046 |
| Final particles (no.) | 98,358 | 98,358 | 98,640 | 28,046 |
| Point-group or helical symmetry | - | - | - | - |
| Estimated error of translations/rotations | - | - | - | - |
| Resolution (global, Å) | 3.4 | 3.4 | 2.9 | 3.6 |
| FSC 0.5 (unmasked/masked) | - | - | - | - |
| FSC 0.143 (unmasked/masked) | 4/3.4 | 4/3.4 | 3.4/2.9 | 4.3/3.6 |
| Resolution range (local, Å) | 3-7 | 3-7 | 3-7 | 3.5-7 |
| Resolution range due to anisotropy (Å) | - | - | - | - |
| Map sharpening *B* factor (Å<sup>2</sup>) | - | - | - | - |
| Map sharpening methods | - | - | - | - |
| **Model composition (for each model)** |  |  |  |  |
| Protein | 1109 | 1109 | 844 | 1005 |
| Ligands | 0 | 0 | 0 | 0 |
| RNA/DNA | 0 | 0 | 4 | 4 |
| **Model Refinement (for each model)** |  |  |  |  |
| Refinement package | Phenix | Phenix | Phenix | Phenix |
| - real or reciprocal space | Real space | Real space | Real space | Real space |
| - resolution cutoff | 3.3 | 3.3 | 2.9 | 3.6 |
| Model-Map scores |  |  |  |  |
| -CC | 0.84 (mask) | 0.85 (mask) | 0.85 (mask) | 0.73 (mask) |
| - FSC (d FSC model at 0.143) | 3.3 (mask) | 3.3 (mask) | 2.9 (mask) | 3.5 (mask) |
| B factors (Å<sup>2</sup>) |  |  |  |  |
| Protein residues | 142.45 | 142.67 | 141.11 | 99.96 |
| Ligands | - | - | - | - |
| RNA/DNA | - | - | 112.56 | 51.93 |
| R.m.s. deviations from ideal values |  |  |  |  |
| Bond lengths (Å) | 0.003 | 0.002 | 0.005 | 0.003 |
| Bond angles (°) | 0.527 | 0.513 | 0.757 | 0.637 |
| **Validation (for each model)** |  |  |  |  |
| MolProbity score | 1.56 | 1.57 | 2.48 | 1.88 |
| CaBLAM outliers | 0.74 | 0.93 | 2.96 | 0.65 |
| Clashscore | 10.92 | 11.46 | 13.57 | 15.78 |
| Poor rotamers (%) | 0.48 | 0.48 | 6.57 | 0.96 |
| C-beta deviations | 0 | 0 | 0 | 0 |
| EMRinger score | - | - | - | - |
| Ramachandran plot |  |  |  |  |
| Favored (%) | 98.17 | 98.26 | 96.62 | 96.99 |
| Outliers (%) | 0.37 | 0.37 | 0.00 | 0.21 |

## Notes

### Summary of Updates

New data was added which included: – a phylogenetic analysis of Cap1 homologs – demonstration that Cap1 also mediated immunity against phage lambda in E. coli, a Gram-negative organism

